# The PPR-related splicing cofactor MSP1/EMB1025 protein, encoded by At4g20090, encode an essential protein that is required for the splicing of *nad1* intron 1 and for the biogenesis of complex I in Arabidopsis mitochondria

**DOI:** 10.1101/615856

**Authors:** Corinne Best, Michal Zmudjak, Oren Ostersetzer-Biran

## Abstract

Group II introns are particularly plentiful within plant mitochondrial genomes (mtDNAs), where they interrupt the coding-regions of many organellar genes, especialy within complex I (CI) subunits. Their splicing is essential for the biogenesis of the respiratory system and is facilitated by various protein-cofactors that belong to a diverse set of RNA-binding cofactors. These including maturases, which co-evolved with their host-introns, and various *trans*-acting factors, such as members of the pentatricopeptide-repeat (PPR) protein family. The genomes of angiosperms contain hundreds of *PPR*-related genes that are postulated to reside within the organelles and affect diverse posttranscriptional steps, such as editing, RNA-stability and processing or translation. Here, we report the characterization of MSP1 (Mitochondria Splicing PPR-factor 1; also denoted as EMB1025), which plays a key role in the processing of *nad1* pre-RNAs in Arabidopsis mitochondria. Mutations in *MSP1* gene-locus (At4g20090) result in early embryonic arrest. To analyze the putative roles of MSP1 in organellar RNA-metabolism we used a modified embryo-rescue method, which allowed us to obtain sufficient plant tissue for the analysis of the RNA and protein profiles associated with *msp1* mutants. Our data indicate that MSP1 is essential for the *trans*-splicing of *nad1* intron 1 in Arabidopsis mitochondria. Accordingly, *msp1* mutants show CI biogenesis defects and reduced respiratory-mediated functions. These results provide with important insights into the roles of nuclear-encoded factors during early plant development, and contribute to our limited understanding of the importance of RNA-maturation and splicing in plant mitochondria during early embryogenesis.

## Introduction

Plants are able to coordinate their energy demands during particular growth and developmental stages, by utilizing complex signaling pathways between their nuclear and the organellar genomes (reviewed by *e.g*., (Gualberto and Kuhn 2014, Gualberto and Newton 2017). Although mitochondria contain their own genomes (mtDNAs), they encode only a few structural RNAs and proteins, whereas the majority of the organellar proteins are encoded by nuclear loci (Huang *et al*. 2013, Jacoby *et al*. 2012, Lee *et al*. 2013, Millar *et al*. 2011). In fact, the organellar ribosomes and the respiratory complexes are assemblies of proteins that are synthesized by the two remote genomes machineries (see *e.g.* (Huang *et al*. 2014, Schertl and Braun 2014). This necessitates complex regulatory mechanisms to allow the stoichiometric accumulation of subunits encoded by both nuclear and organellar genomes (Woodson and Chory 2008).

Accumulating data indicate that the regulation of RNA-processing plays a pivotal role in the expression of organellar genes in plants (Brown *et al*. 2014, Gualberto and Kuhn 2014, Hammani and Giege 2014, Liere *et al*. 2011, Schmitz-Linneweber *et al*. 2015, Small 2013, Small *et al*. 2013). The significance of posttranscriptional regulation in plant mitochondria is further reflected by the extended half-lives of many mtRNAs, and the fact that their translation seemed uncoupled from their transcription (Kühn *et al*. 2007, Kühn *et al*. 2005, Liere and Börner 2011, Liere, *et al*. 2011, Planchard *et al*. 2018, Zmudjak and Ostersetzer-Biran 2017). Accordingly, altered mtRNA metabolism often result with severe growth and developmental defect phenotypes.

One of the most remarkable features of plant mitochondria is the presence of intervening sequences, which reside in the coding-regions of many of the organellar genes. RNA-splicing, which removes the introns from the pre-mRNAs, is a critical step in the posttranscriptional regulation of plant organellar gene-expression and relies upon the activities of various protein cofactors (Brown, *et al*. 2014, Hammani and Giege 2014, Liere, *et al*. 2011, Schmitz-Linneweber, *et al*. 2015, Small 2013, Zmudjak and Ostersetzer-Biran 2017, Zmudjak *et al*. 2017). These belong to a diverse set of RNA-binding protein families, which are encoded (primarily) in the nucleus and post-translationary imported into the mitochondrion (Zmudjak and Ostersetzer-Biran 2017).

The introns found within the mtDNAs of land-plants are generally classified as group II-type sequences that interrupt the coding sequences of protein-encoding genes, and are found predominantly within various complex I subunits (Bonen 2008, Brown, *et al*. 2014, Schmitz-Linneweber, *et al*. 2015). Based on structural similarities, their catalytic activities (*i.e.*, two trans-esterification reactions leading to the splicing of the host exons and the release of the intron in a lariat form) and retrohoming ability, it is postulated that these introns are the progenitors both the spliceosomal introns and transposons (Cech 1986). Although few canonical introns in this class are able to catalyze their own excision *in vitro* (i.e., self-splicing, autocatalytic intron RNAs), the splicing of virtually all group introns *in vivo* relies on the activities of various cofactors. In bacteria and yeast mitochondria these involve proteins that are encoded within the introns themselves, known as maturases (Matsuura *et al*. 2001, Qu *et al*. 2016, Schmitz-Linneweber, *et al*. 2015, Zhao and Pyle 2016, Zimmerly *et al*. 2001).

Although the organellar introns in plants have evolved from maturase-encoding group II introns, these have diverged considerably, such as they lack many sequence elements that are considered to be essential for the splicing activity, and also lost the vast majority of their cognate maturase-ORFs (Bonen 2008, Brown, *et al*. 2014, Hanley and Schuler 1988, Zimmerly and Semper 2015). Some became fragmented so they are scattered in the mtDNA, transcribed separately and then spliced in ‘*trans*’. In Arabidopsis, *trans*-spliced introns include the first and third introns in NADH-dehydrogenase subunit 1 (*i.e*. *nad1* introns 1 and 3), *nad2* i2, and *nad5* i2 and i3 (Unseld *et al*. 1997). Due to their degenerate nature and the fact that the mitochondrial introns have lost their evolutionary-related maturases, the splicing of group II introns in plant mitochondrial relies on the activities of various host-encoded proteins (Bonen 2008, Brown, *et al*. 2014, Schmitz-Linneweber, *et al*. 2015).

Genetic (forward and reverse) screens led to the identification of many splicing cofactors in land-plants mitochondria (Brown, *et al*. 2014, Colas des Francs-Small and Small 2014, Hammani and Giege 2014, Schmitz-Linneweber, *et al*. 2015, Zmudjak and Ostersetzer-Biran 2017). These include proteins related to group II maturases (*i.e*. MatR encoded by *nad1* i4 within the mitochondria, and four nuclear-encoded maturase proteins, denoted as nMATs 1-to-4) (Cohen *et al*. 2014, Keren *et al*. 2009, Keren *et al*. 2012, Mohr and Lambowitz 2003, Nakagawa and Sakurai 2006, Schmitz-Linneweber, *et al*. 2015, Sultan *et al*. 2016, Zmudjak, *et al*. 2017). In addition to maturases, proteins that were shown to function in the splicing of group II introns include RNA helicases (He *et al*. 2012, Köhler *et al*. 2010, Matthes *et al*. 2007, Putnam and Jankowsky 2013, Zmudjak, *et al*. 2017), PORR-related factors (Colas des Francs-Small *et al*. 2012) and many proteins belonging to the pentatricopeptide repeat (PPR) family (Colas des Francs-Small *et al*. 2014, Colas des Francs-Small and Small 2014, Falcon de Longevialle *et al*. 2007, Gualberto *et al*. 2015, Koprivova *et al*. 2010, Liu *et al*. 2010, Schmitz-Linneweber, *et al*. 2015, Wang *et al*. 2017, Wang *et al*. 2018, Weissenberger *et al*. 2017, Yang *et al*. 2014, Zmudjak and Ostersetzer-Biran 2017). However, there is still little mechanistic information regarding how this proteins facilitate the splicing of their group II intron RNA-targets, and even less is known about how *trans*-splicing is mediated *in vivo*. The current manuscripts focuses on At4g20090, which encodes a PPR protein that is key to the *trans*-splicing of *nad1* intron 1 in Arabidopsis mitochondria.

Plants encode numerous proteins harboring a variety of RNA binding domains, many of which are predicted to reside in the mitochondria or chloroplasts. Many of these are identified as PPR proteins, which are recognized by the presence of tandem repeats (2 to 30 repeats) of a degenerate 35 amino acid domain termed as the PPR motif (Peeters and Small 2001). Each PPR motif forms two α-helices, which interact with each other and fold into two antiparallel helices (helix A and helix B) connected by a coil (Coquille *et al*. 2014, Gully *et al*. 2015, Kaushal *et al*. 2014, Shen *et al*. 2015, Shen *et al*. 2016). PPRs comprises one of the largest gene-family in flowering plants, with hundreds of genes identified in the complete genomes of various angiosperms (i.e., approximately 450 *PPR* genes in Arabidopsis and about 480 in rice) (Chen *et al*. 2018, Gorchs Rovira and Smith 2019, Lurin *et al*. 2004, Schmitz-Linneweber and Small 2008, Shikanai and Fujii 2013). Based on the number of the PPR repeats and their topologies, the PPRs in plants are divided into two subclasses, teremed as P and PLS. Members of the PLS-subclass contain extra domains (*i.e*., E, E^+^ and the DYW motif), which are absent from the P-subclass proteins that typically consist of only tandem arrays of the conserved PPR motif (Brehme *et al*. 2014, Fujii and Small 2011, Lurin, *et al*. 2004, O’Toole *et al*. 2008, Schmitz-Linneweber and Small 2008, Takenaka *et al*. 2008). The vast majority of the PPRs are predicted to reside within mitochondria or plastids (based on their N’-terminal regions which show homology to organellar targeting presequences), where they are expected to function in various RNA-processing event, such as editing, RNA stability and processing and translation (for recent reviews see *e.g.*, (Gorchs Rovira and Smith 2019, Manna 2015, Shikanai and Fujii 2013, Zmudjak and Ostersetzer-Biran 2017). Many seem to act in a sequence specific manner. Recent analyses designed to decipher the mechanism of RNA-recognition by PPR proteins indicate a similar RNA recognition code to that of the transcription activator-like effectors (TALEs) (Barkan *et al*. 2012, Yagi *et al*. 2013, Yagi *et al*. 2014).

Here, we examined the functions of the P-subclass Mitochondria Splicing PPR-factor 1 (MSP1), encoded by the At4g20090 gene-locus (also annotated as EMBRYO DEFECTIVE 1025, EMB1025), in the metabolism of mitochondrial RNAs (mtRNAs) in Arabidopsis. The effects of lowering the expression of MSP1 on mitochondria functions and RNA metabolism and the physiology of *msp1* knockout-lines in Arabidopsis are discussed.

## Results

### The topology of MSP1 protein

PPR proteins are present in most eukaryotes, but have undergone a huge expansion in the genomes of land-plants (Hayes *et al*. 2012). Many of these are identified as sequence-specific RNA-binding cofactors that are involved in various RNA processing events (Dahan and Mireau 2013, Gorchs Rovira and Smith 2019, Manna 2015, Pfalz *et al*. 2009, Ruwe and Schmitz-Linneweber 2011). The Arabidopsis SeedGenes database (Meinke *et al*. 2008) contains hundreds of genes (with about 900 different mutant lines) that were found to be required during embryo development. Many of the *EMBRYO-DEFECTIVE* (*EMB*) genes encode proteins with an essential function required throughout the life cycle (Muralla *et al*. 2011). Among these, 32 are identified as PPR-related proteins (http://seedgenes.org/GeneList.html), including OTP43 encoded by At1g74900 that was shown to influence the maturation of *nad1* i1 (Falcon de Longevialle, *et al*. 2007). Here, we focus on At4g20090, which is annotated as EMBRYO DEFECTIVE 1025 (EMB1025; SeedGenes database). We altered the annotation of EMB1025 into MSP1 (Mitochondria Splicing PPR-factor 1) to better describe the molecular functions of At4g20090 gene-product in mitochondrial RNA-processing (see below).

Domain search analysis of the Arabidopsis MSP1 protein (660 aa), using the SMART (Letunic *et al*. 2012) and Conserved Domain Database (CDD) (Marchler-Bauer *et al*. 2003) servers indicated the presence of 15 PPR motifs, with a predicted topology of: *NH_2_*-76-P-P-1-P-4-P-P-P-P-P-P-P-P-P-P-3-P-P-50-*COOH*, in which ‘P’ indicate to canonical PPR motifs (35-36 aa), while amino acids not assigned to any defined domain are indicated by numbers (Fig. 1a, Suppl. Fig. S1a and Table S1). Mitochondrial presequence cleavage site prediction servers, available at the ExPASy portal (https://www.expasy.org), indicate a short N-terminal region (*i.e.*, 36-37 amino acids, Fig. S1a, underlined) typical to organellar targeting regions which may be processed by either the Mitochondrial Processing Peptidase (MPP, 36 aa) or Intermediate Cleavage Peptidase 55 (ICP55, 37 aa) (Figs. 1 and S1a).

**Figure 1.**
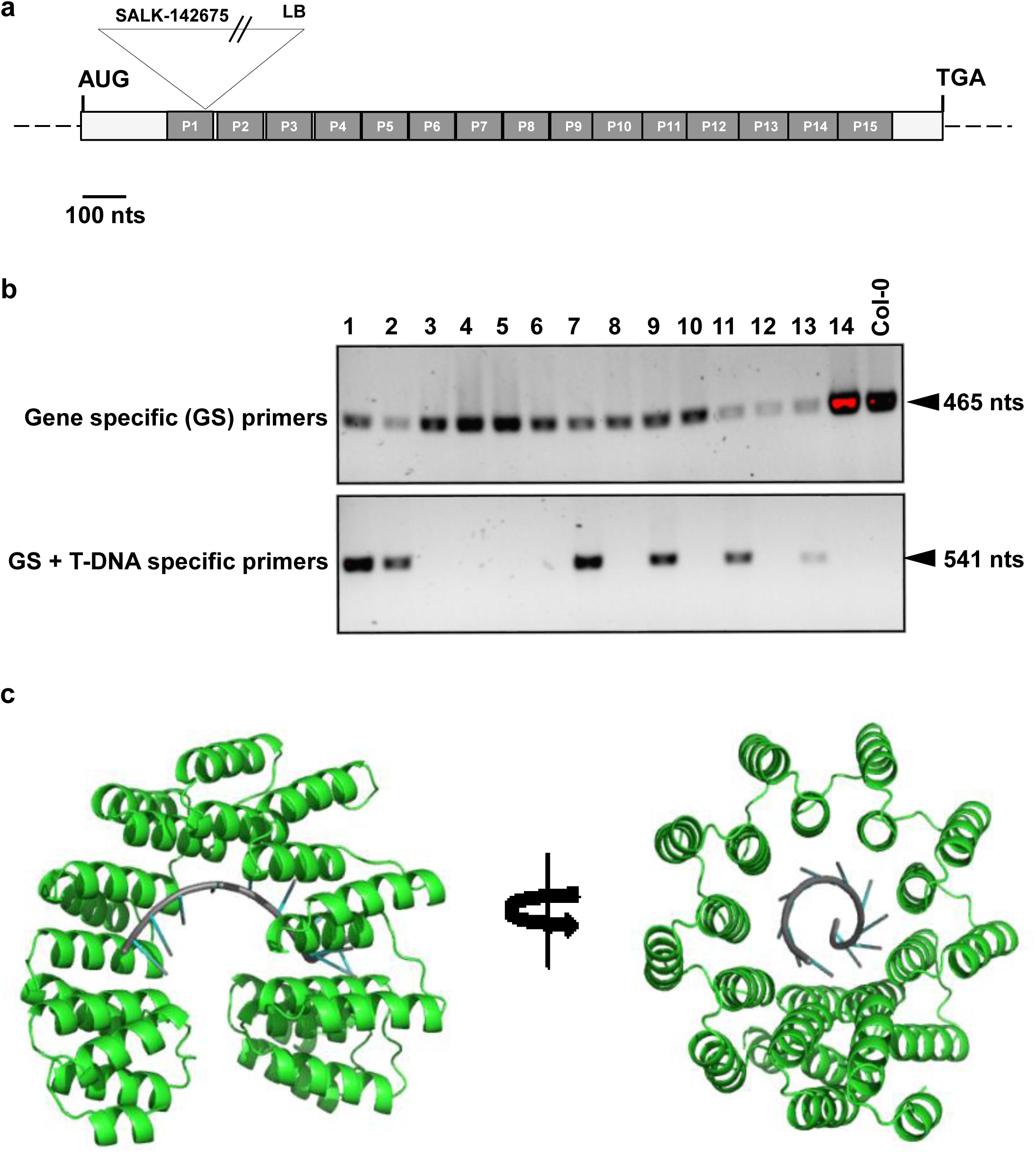
MSP1/EMB1025 (At4g20090) gene and protein topologies. (a) Scheme of At4g20090 gene structure. In silico analysis indicate the presence of 15 repeats of the conserved PPR motif (see also Supplementary Fig. S1 and Table S1). The position of the T-DNA insertion site in the coding region of MSP1 (i.e., SALK-line 142675, *msp1-1*) is found 312 nucleotides downstream to the first ATG, within the first PPR motif, as indicated established by PCR and sequencing. (b) PCR screening of soil-germinated plantlets obtained from mature seeds of heterozygous *msp1-1* mutants, with primers designed to the T-DNA (LBb1.3) and *MSP1* gene (see Table SX). Arrows indicate 500 nucleotides. (c) Homology modeling of the *Arabidopsis thaliana* MSP1 protein. To get more insights into MSP1 mode of action and RNA recognition, we performed an atomic 3D model of MSP1 protein, using the Phyre^2^ (Kelley and Sternberg 2009) and ROBETTA (Kim, *et al*. 2004) servers. A hypothetical 3D structure of MSP1 protein was visualized by the PyMol software.

Analysis of the predicted sequence of MSP1 protein, using the Phyre^2^ (Kelley and Sternberg 2009) and ROBETTA (Kim *et al*. 2004) servers, indicated that At-MSP1 shares a similar fold (Fig. 1c and Suppl. Fig. S1b) with serval PPR proteins, such as the PPR10 protein from maize (Gully, *et al*. 2015), the mammalian 28S ribosomal protein S5 (MRPS5) (Kaushal, *et al*. 2014) or the synthetically-designed dPPR-U10 (Shen, *et al*. 2015, Shen, *et al*. 2016) and cPPR-polyC proteins (Coquille, *et al*. 2014). The predicted 3D structure of At-MSP1 predicts a right-handed two-turn superhelical fold (Gully, *et al*. 2015, Yin *et al*. 2013), with 15 PPR motifs (amino acids 76-609) surrounded by 4 short α-helices at the N’-terminal region and three short α-helices at the C-terminus (Suppl. Fig. S1b). These amino-terminal regions may contribute to ligand specificity (Gully, *et al*. 2015, Yin, *et al*. 2013), as also shown in the cases of the repeat-domains TPR and TALE protein families (Deng *et al*. 2012, Grove *et al*. 2008). We further speculated that the internal core of MSP1 protein, which contains predicted basic regions (Suppl. Fig. S1b, labeled in blue color), may correspond to RNA-binding (Fig. 1c), as indicated for the PPR10 protein (Gully, *et al*. 2015, Yin, *et al*. 2013). However, additional structural studies and biochemical assays would be required to confirm the hypothetical 3D model of MSP1 protein bound to RNAs (Fig. 1c), and to analyze its binding characteristics.

### MSP1 is located within the mitochondria *in vivo*

Based on its N-terminal region, MSP1 is predicted to be localized to the chloroplasts or mitochondria in Arabidopsis (Hooper *et al*. 2017). MSP1 could not be found among the different proteins identified in mass-spectrometry analyses of plant organellar fractions, which include many other proteins with predicted RNA-mediated functions (including several PPR proteins). Genome-wide analysis and global localization assays of Arabidopsis PPR proteins with green-(GFP) and red-fluorescent (RFP) protein fusions, suggested that At4g20090, among various other PPR proteins in Arabidopsis analyzed therein, resides within the mitochondria (Lurin, *et al*. 2004). As bioinformatics tools still remain inaccurate, ascribing different proteins to the wrong intracellular site(s) (Heazlewood *et al*. 2005), and the global nature of previous localization analyses showing that MSP1 may reside within the mitochondria (Lurin, *et al*. 2004), the localization of MSP1 needs to be further supported experimentally. For this purpose, constructs encoding the N’-terminal region (150 amino acids) of *MSP1* gene (*i.e*., *N’*-MSP1-GFP) in Arabidopsis were cloned in-frame to GFP, introduced into tobacco plants, and the location of the GFP-fusion protein was determined by confocal microscopy (Fig. 2a). While GFP alone showed cytosolic distribution of the GFP signal, with considerable diffusion into the nucleus, the signal of the *N’*-MSP1-GFP fusion protein was detected as rod-shaped granules co-localizing with those of MitoTracker, a mitochondrion-specific fluorescent probe (Fig. 2b). These results coincide with the protein localization predictions and the data in (Lurin, *et al*. 2004), which together strongly support the localization of MSP1 to the mitochondria in angiosperms.

**Fig 2.**
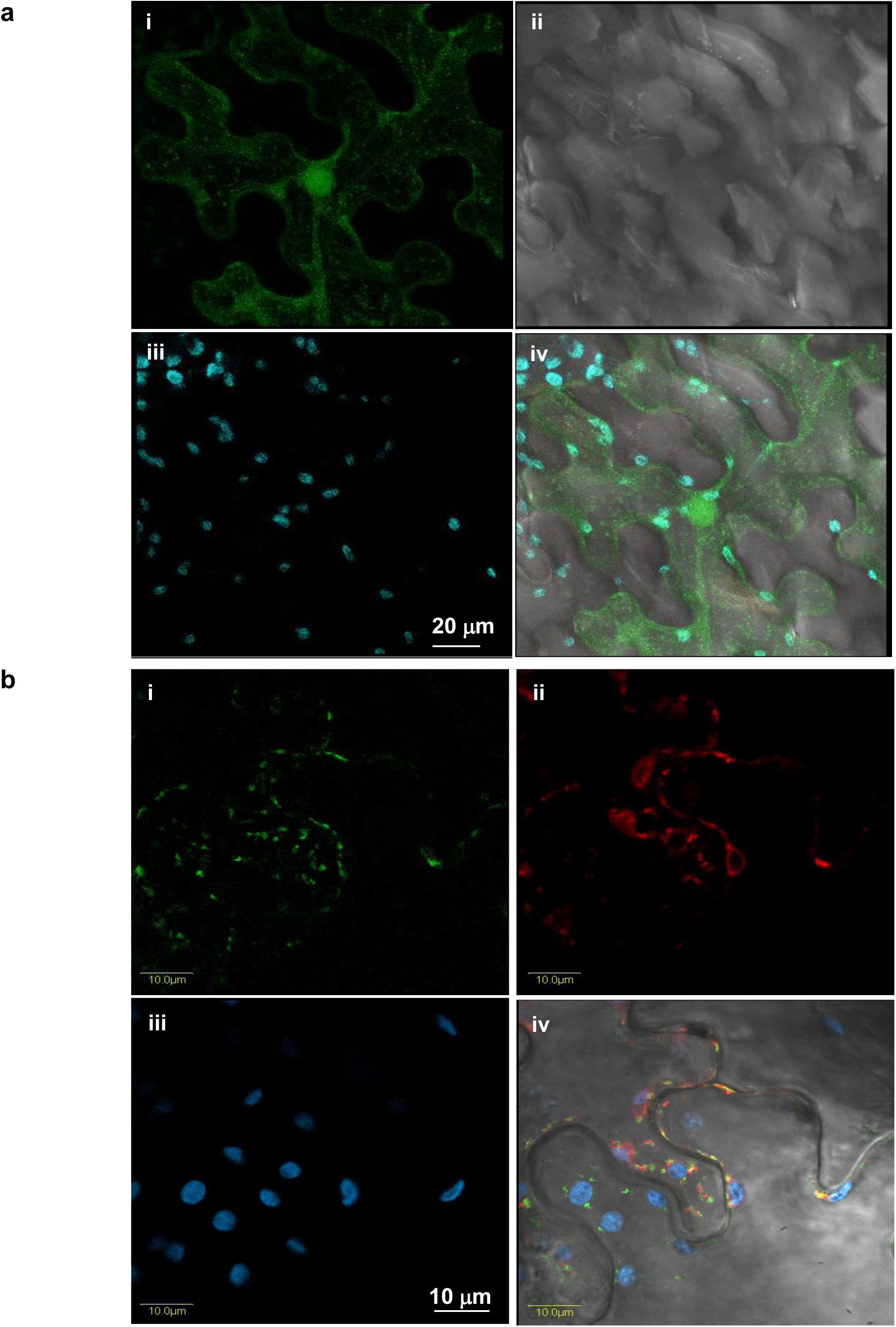
GFP localization assays indicate that MSP1 is localized to the mitochondria, in vivo. Tobacco leaves were agroinfiltrated with GFP alone (a) or GFP fused in frame to the N-terminal region (150 amino acids) of the Arabidopsis MSP1 (At4g20090) protein (b). The GFP signals (green, upper left a*i* and b*i*), lower-epidermis image (grey color, upper right panel a*ii*), MitoTracker marker (red, upper right panel b*ii*), chlorophyll autofluorescence (blue, lower left panels a*iii* and b*iii*) and merged images (lower right panels a*iv* and b*iv*).

### The Arabidopsis *MSP1* gene-locus (At4g20090) encodes an essential protein that is required during early embryogenesis

For expression analysis, we used microarray and high throughput sequencing experiments available in the public domain. These include ‘The Arabidopsis Information Resource’ (TAIR; http://www.arabidopsis.org) database and the Genevestigator analysis toolbox (Hruz *et al*. 2008, Zimmermann *et al*. 2004). The expression data indicated differential expression of At4g20090 through development in different organs, where MSP1 exhibits dominant expression in embryonic organs, young developing leaves, apical root tissues, flowers, the shoot apex (Fig. S2). These databases indicated that MSP1 accumulates to high levels particularly during imbibition and early germination stages (Fig. S2).

To analyze the putative roles of MSP1 in plant mitochondrial biogenesis and function, we screened available T-DNA lines for insertions in the At4g20090 gene-locus in Arabidopsis, including SALK-142675 (*msp1-1*), SALK-018927 (*msp1-2*) and SALK-070654 (*msp1-3*). Sequencing of genomic PCR products, spanning the T-DNA insertion junction in heterozygous *msp1-1* mutant-plants (SALK-142675 line), confirmed that the T-DNA was inserted within the coding region of the intronless *MSP1* gene sequence (Fig. 1a and Suppl. Fig. S1). Homozygous SALK-018927 and SALK-070654 plants (containing T-DNA insertions at the 5’ or 3’ termini of the *MSP1* gene, respectively) did not show any obvious phenotypes under normal growth conditions (see Material and Methods). Plants homozygous for the T-DNA insertions at both these loci were identified among the analyzed population, but RT-qPCR analysis indicated that both these T-DNA insertions did not altered the expression of MSP1 in the homozygous lines.

However, no plants homozygous for the *msp1-1* mutant allele could be recovered among the seeds obtained from heterozygote SALK-142675 line. The selfed progeny of the heterozygous *msp1-1* mutants showed a kanamycin resistant/sensitive segregation ratio of about 2:1 (*i.e.*, 119 out of 175 seedlings), suggesting that At4g20090 encodes an essential gene. PCR genotyping further indicated that the resistant lines were all heterozygous for this mutation (Fig. 1b and Suppl. Fig. S3). The heterozygous SALK-142675 plants were phenotypically indistinguishable from the wild-type plants (Col-0), suggesting that the T-DNA insertion within the coding region of *MSP1* gene result in a recessive embryo-lethal phenotype and that a single copy of the gene is sufficient to support normal growth and development.

To confirm this assumption, we compared the embryo-development in seeds of immature siliques (about 4 to 6 days following fertilization) of heterozygous *msp1* plants (Fig. 3a). About quarter of the progeny in young immature siliques of SALK-142675 line (*i.e*., 25.1 ± 4.3% of 287 seeds examined under the microscope) contained white-translucent seeds (Fig. 3b). During the maturation of the siliques, the white seeds either collapsed into seeds lacking embryos (Fig. 3a, indicated by “1”), or degenerated into shrunken brown seeds, arrested at the globular stage (Fig. 3a, indicated by “2”). Genotyping by PCR indicated that these were all homozygous for the loss-of-function allele. Similar to *msp1-1*, a variation between homozygous individuals within loss-of-function alleles, where also observed in the cases of *ndufv1* (Kuhn *et al*. 2015) and *cod1* (Dahan *et al*. 2014) mutants that are affected in the biogenesis of the mitochondrial complexes I or III, respectively. This phenomenon may relate to some differences in the production, storage or mobilization of reserves stored in the endosperm of the developing seeds (Focks and Benning 1998, Keren, *et al*. 2012, Kuhn, *et al*. 2015, Pinfield-Wells *et al*. 2005). Nomarski microscopy analyses indicated that green seeds (*i.e.*, wild-type or heterozygous seeds) in the young siliques carried mature embryos (Figs. 3b and 3c, labeled as “i”), while the white seeds of the same progeny showed delayed development with embryos which were arrested in the early globular stage (Figs. 3c, labeled as “ii”) (http://arabidopsisbook.org).

**Figure 3.**
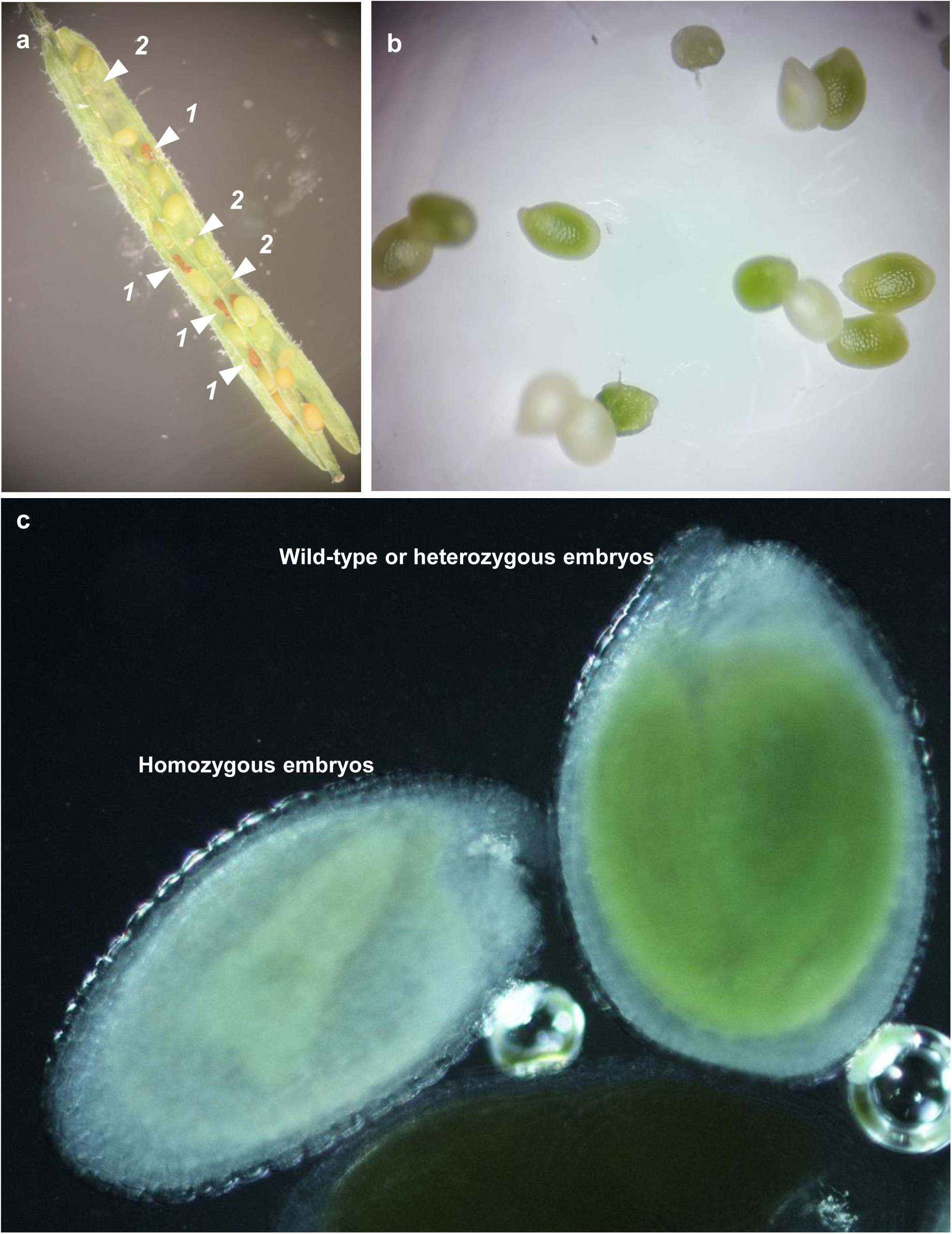
Knockout *msp1-1* mutants show embryo arrested phenotypes. Mature siliques obtained from heterozygous *msp1-1* (SALK-142675) lines (a). Arrows indicate homozygous mutant seeds. Numbers indicate to seeds which contain embryos arrested at the late heart or torpedo stages, while ‘2’ show aborted seeds with dead embryos. (b) White (containing homozygous embryos) and green seeds (contacting either heterozygous or wild-type embryos) obtained from immature siliques of *msp1-1*. (c) Images taken with differential interference contrast (*i.e.*, Nomarski) microscopy.

### A modified method for embryo rescue of Arabidopsis *msp1-1* mutant-lines

Some Arabidopsis mutants impaired in mitochondria functions seem to be unable to germinate under standard conditions, due to defects in embryo development (see *e.g.*, (Colas des Francs-Small and Small 2014, Cordoba *et al*. 2016, Dahan, *et al*. 2014, Focks and Benning 1998, Franzmann *et al*. 1989, Fromm *et al*. 2016a, Keren, *et al*. 2009, Kuhn, *et al*. 2015, Ostersetzer-Biran 2016, Pinfield-Wells, *et al*. 2005). As loss-of-function knockout *msp1* mutant-seeds were not informative, we developed an optimized embryo-rescue strategy that has been modified from the methods described by Franzmann, *et al*. (1989) and recent approaches used to recover germination-arrested phenotypes in Arabidopsis plants affected in mitochondria biogenesis, as the PPR-related *cod1* mutant (Dahan, *et al*. 2014) and *cal1/cal2* mutants (Cordoba, *et al*. 2016, Fromm, *et al*. 2016a).

As previously described for *cod1* and *cal1/cal2*, the matured dried seeds of *msp1* mutants were unable to germinate on soil under standard conditions. However, the settings described to successfully rescue *cod1* (Dahan, *et al*. 2014) or *cal1/cal2* mutants (Fromm, *et al*. 2016a), where found as unsuitable for the germination of mature or immature (white) *msp1* seeds, even when transferred to MS-agar plates supplemented with different concentrations of sugar in the medium (*i.e*., 1-5% sucrose). These results may relate to the embryogenesis defect phenotypes of *msp1-1* mutants that are arrested at the early torpedo stage (Fig. 3), whereas the seeds of *cod1* and *cal1/cal2* seem to contain small but otherwise fully developed embryos (Dahan, *et al*. 2014).

The optimized conditions for germination of homozygous knockout *msp1* mutant-allele was obtained when white seeds, containing homozygous embryos, were selected from sterilized immature siliques of heterozygous *msp1-1* plants, were sown on MS-agar plates supplemented with 1 to 3% sucrose and various vitamins and then transferred to growth chamber (AR-41L3, Percival) and kept under standard growth conditions (see Materials and Methods). Under the *in vitro* conditions, a small number of the cultivated *msp1* seeds (*i.e*., about 1∼2%) germinated following 3 weeks in the growth chamber, and about 30% of the seeds were able to germinate following 3 months of incubation in the growth chamber (Fig. 4a). The germinated plantlets slowly (3 to 4 months) developed into a 6 to 8 leaf-stage (stage R6) (Boyes *et al*. 2001), but were unable to continue their development beyond this stage on the MS-agar plates (Fig. 4a). Longer incubation times on the MS-plates resulted with bleaching and death of the plantlets. For DNA and RNA analyses, we used 3-month-old seedlings (staring initial imbibition), arrested at the 6∼8 leaf stage.

**Figure 4.**
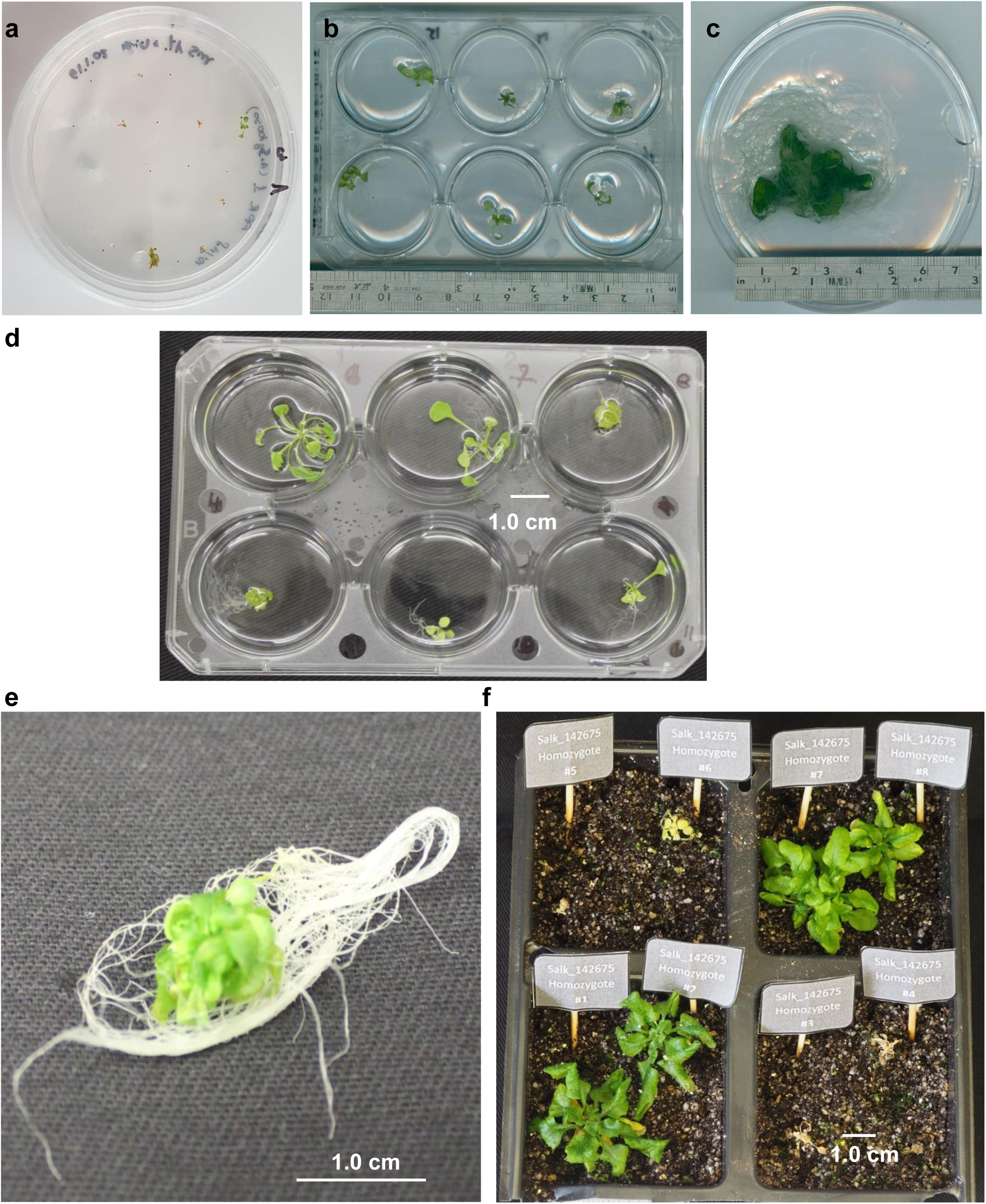
Plant phenotypes associated with rescued homozygous *msp1-1* mutant line. Seeds were collected under sterile conditions from surface sterilized immature siliques of Col-O and heterozygous *msp1-1* (SALK-line 142675) plants. (a) White homozygous seeds, obtained from immature siliques of *msp1-1* mutant-plants, were sown on petri-dishes containing MS-Agar supplemented with 1% sucrose and various vitamins (see Materials and Methods) and grown in Arabidopsis growth chamber under standard conditions. 6-week-old plantlets grown the MS-plates, obtained from white seeds (b) and immature wild-type seeds at the torpedo stage (c), were transferred to liquid MS media supplemented with 1% sucrose and vitamins (see Materials and Methods). (d) Images of 4-months old rescued *msp1* and wild-type seedlings showing variability among individual rescued *msp1-1* plantlets. (e) While some of the seedlings develop a typical rosette, some of the plantlets develop short leaves with limited stems, leading to a bushy-like phenotype (Dahan, *et al*. 2014). (f) 5 month-old rosettes from homozygous *msp1-1* plants growing on soil in short-day conditions.

Remarkably, when the *msp1* plantlets where transplanted into a liquid MS-based medium containing 1% sucrose and vitamins (see Materials and Methods) their development exceed beyond the R6 stage, where significant increases in both leaf and root biomass were observed following 2∼3 weeks of incubation in the MS-media (Fig. 4b-4e, and Materials and Methods). Under these conditions, the growth rates of Arabidopsis wild-type seedlings was also notably increased during the cultivation time (Fig. 4c).

A variation between individuals within the rescued *msp1-1* line was also visible. While some seedlings developed a typical rosette, others exhibited a bushy structure with miniature leaves morphology (Figs. 4d and 4e), an abnormal developmental phenotype that was previously reported for some rescued *emb* mutants (Franzmann, *et al*. 1989) and *cod1* (Dahan, *et al*. 2014). When the liquid-grown plantlets where transferred to the soil, the subsequent growth of *msp1-1* plants was retarded (Fig. 4f), as compared with immature wild-type seeds rescued at the torpedo stage. These include small plants with a curly-leaf morphology, which was very similar to that of *opt43* (Colas des Francs-Small and Small 2014) and other Arabidopsis mutants affected in mitochondria biogenesis. None of the rescued *msp1-1* plants where able to set flowers or to produce viable seeds.

### Mutations in the *MPS1* gene-loci strongly affect the accumulation of mature *nad1* transcripts in Arabidopsis mitochondria

The homology of MSP1 with organellar P-subclass PPR proteins and its localization to the mitochondria in plants support a role for this protein in the processing of organellar transcripts in Arabidopsis plants. To gain more insights into the roles of MSP1 in mitochondria RNA (*i.e*., mtRNAs) metabolism, and to analyze the effects of mutations in *MSP1* gene on the steady-state levels of the different transcripts in Arabidopsis mitochondria, we used transcriptome analyses by quantitative reverse transcription PCRs (see *e.g*. (Cohen, *et al*. 2014, Colas des Francs-Small, *et al*. 2012, Keren, *et al*. 2012, Sultan, *et al*. 2016, Zmudjak *et al*. 2013). The relative accumulation of organellar transcripts was evaluated from the relative ratios of various mRNAs in embryo-rescued *msp1-1* (Fig. 5a) or wild-type (Fig. 5b) plantlets versus those observed in 3-week-old MS-agar grown wild-type plants.

**Figure 5.**
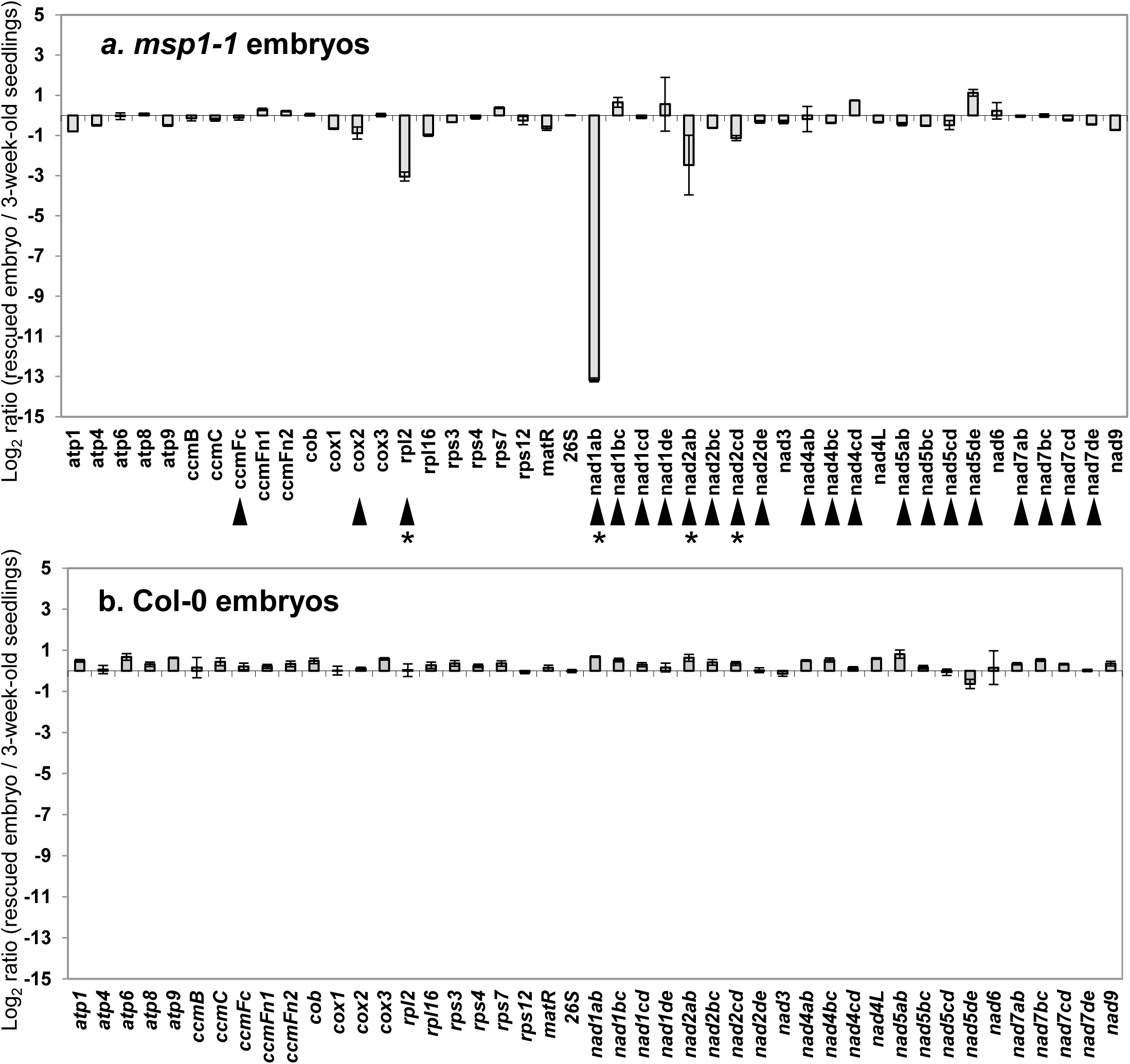
Transcript abundance of mitochondrial mRNAs in *msp1-1* mutants. Transcriptome analyses of mitochondria mRNAs levels in Arabidopsis plants by RT-qPCR was preformed essentially as described previously (Cohen, *et al*. 2014, Sultan, *et al*. 2016, Zmudjak, *et al*. 2013). RNA extracted from 3-week-old seedlings of wild-type (Col-0) plants, rescued *msp1* mutants and plantlets germinated from seeds obtained from immature siliques of Col-0 plants at the torpedo stage (Fig. 3), was reverse-transcribed, and the relative steady-state levels of cDNAs corresponding to the different organellar transcripts were evaluated by qPCR with primers which specifically amplified mRNAs (Table S3). The histogram shows the relative mRNAs levels (*i.e*. log2 ratios) in *msp1* mutant line (a) and rescued Col-0 plantlets (b) versus those of wild-type plants. Transcripts analyzed in these assays include the NADH dehydrogenase (CI) subunits *nad1* exons a-b, b-c, c-d, d-e, *nad2* exons a-b, b-c, c-d, d-e, *nad3*, *nad4* exons a-b, b-c, c-d, *nad4L*, *nad5* exons a-b, b-c, c-d, d-e, the intronless *nad6* subunit, *nad7* exons a-b, b-c, c-d, d-e, and *nad9*, the complex III *cob* subunit, cytochrome oxidase (complex IV) *cox1*, *cox2* exons a-b and *cox3* subunits, the ATP synthase (i.e., complex V) subunits *atp1*, *atp4*, *atp6*, *atp8* and *atp9*, genes encoding the cytochrome c maturation (*ccm*) factors, *ccmB*, *ccmC*, *ccmFn1*, *ccmFn2* and *ccmFc* exons a-b, the ribosomal subunits *rpl2* exons a-b, *rps3* exons a-b, *rps4*, *rps7*, *rps12*, *rpl16*, *rrn26*, and the *mttB* gene. Arrows indicate to genes that are interrupted by group II intron sequences in Arabidopsis mitochondria (Unseld, *et al*. 1997), while asterisks indicate to transcripts where the mRNA levels were reduced in the *msp1-1* line. The values are means of three biological replicates (error bars indicate one standard deviation).

A notable reduction in the accumulation of mtRNA was observed in the cases of mature transcripts corresponding to the complex I subunit *nad1* exons a-b (*nad1ab*), in which the calculated mRNA levels in *msp1-1* mutant was about 10,000-fold (*i.e.*, ∼2^13) lower than that in the wild-type plants (Fig. 5a). We also noticed to milder reductions in the steady-state levels of transcripts corresponding to *nad2* exons a-b (5.58-fold), *nad2* exons c-d (2.28-fold) and the ribosomal *rpl2* subunit mRNA (8.28-fold) in msp1-1 mutants (Fig. 5a), which together with *nad1ab* are all containing group II introns in *Arabidopsis* mitochondria (Unseld, *et al*. 1997). The expression of other transcripts, including *ccmFc*, *cox2*, *nad4*, *nad5*, *nad7* and *rps3* genes that their genes are also interrupted by group II introns sequences, was not significantly affected in the rescued *msp1-1* mutant line. Similarly, the accumulation of mRNAs corresponding to various other mitochondrial genes, which lack group II introns (*i.e*. ‘intronless’ transcripts), was not significantly affected by the mutation in *MSP1* gene locus (Fig. 5a). These included *cox1* and *cox3* subunits of complex IV, subunits of the ATP synthase complex (CV), cytochrome C biogenesis and maturation (*ccm*) factors, and various ribosomal genes (other than *rps3*), which their mRNAs levels in *msp1* were comparable to those seen in the wild-type plants (Fig. 5a). The RNA profiles of seedlings obtained from immature seeds of wild-type plants at the globular stage and grown under the same conditions did not show any significant alterations in the accumulation of mitochondrial mRNA transcripts (Fig. 5b).

### MSP1 functions in the *trans*-splicing of *nad1* intron 1 in Arabidopsis mitochondria

Reduced signals corresponding to mature *nad1ab*, *nad2ab*, *nad2cd* and *rpl2* transcripts may indicate to splicing defects in *msp1*, as was previously indicated for various PPR factors involved in organellar group II introns splicing in plants (Colas des Francs-Small, *et al*. 2014, Falcon de Longevialle, *et al*. 2007, Gualberto, *et al*. 2015, Koprivova, *et al*. 2010, Liu, *et al*. 2010, Wang, *et al*. 2017, Wang, *et al*. 2018, Weissenberger, *et al*. 2017, Yang, *et al*. 2014). To further establish the putative roles of MSP1 in the processing of group II introns in Arabidopsis, we compared the splicing efficiencies (*i.e*. the ratios of pre-RNAs to mRNAs in mutants versus wild-type plants) of the 23 introns found in the mitochondrial DNA of *Arabidopsis thaliana* plants (Unseld, *et al*. 1997), between wild-type (Col-0) and *msp1* plants, by RT-qPCRs of total mtRNAs obtained from wild-type plants and the *msp1* mutant-line. Splicing defects were determined to be present only in cases where the accumulation of a specific pre-mRNA was correlated with a reduced level of its corresponding mRNA in the mutant.

The analyses of the mRNA to pre-RNA ratios in *msp1-1* plant versus wild-type indicated an extremely strong defect in the maturation of *nad1* intron 1 (*i.e*., *nad1 i1*, Fig. 6a), where the splicing efficiency for this intron was reduced by about 250,000-fold (*i.e.*, ∼2^18) lower than the wild-type plants. To some extent, reduced splicing efficiency was also observed in the case of *nad2* i1 (*i.e*., about 4x lower) (Fig. 6a). However, the splicing efficiencies of *nad2* i3 and the single intron in *rpl2* were not significantly affected in *msp1-1* mutant-line (Fig. 6a). The splicing efficiencies of the remaining 19 introns in the mitochondria of *msp1-1* mutant were found similar to those of the wild-type plants (Fig. 6a).

**Figure 6.**
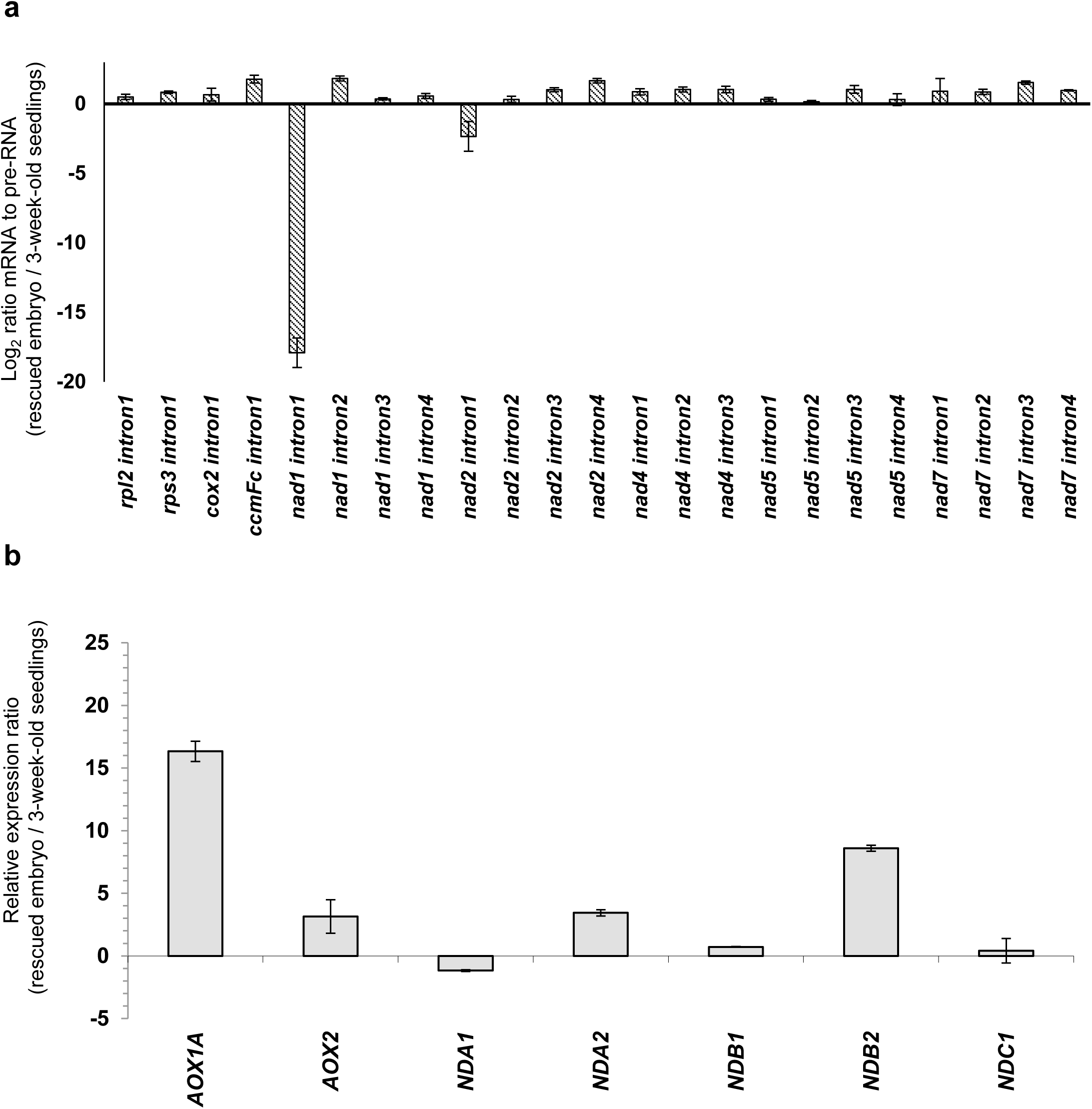
Splicing efficiencies in *msp1* mutants. (a) The relative accumulation of mRNA and pre-RNA transcripts in wild-type and *msp1* plants, corresponding to the 23 group II intron sequences in Arabidopsis, was evaluated by RT-qPCR, essentially as described previously (Cohen, *et al*. 2014, Sultan, *et al*. 2016, Zmudjak, *et al*. 2013). RNA extracted from 3-week-old seedlings of wild-type (Col-0) plants, rescued *msp1* mutants and plantlets germinated from seeds obtained from immature siliques of Col-0 plants at the torpedo stage (Fig. 3), was reverse-transcribed, and the relative steady-state levels of cDNAs corresponding to the different organellar transcripts were evaluated by qPCR with primers which specifically amplified pre-RNAs and mRNAs (Table S3). The histogram shows the splicing efficiencies as indicated by the log2 ratios of pre-RNA to mRNA transcript abundance in *msp1* mutant lines compared with those of wild-type plants. The values are means of three biological replicates (error bars indicate one standard deviation). (b) The relative steady-state levels of mRNAs corresponding to various AOXs and alternative NAD(P)H isoforms was determined by RT-qPCR in 3 week-old wild-type and embryo-rescued *msp1* plants after normalization to the actin2 (At3g1878) and 18S rRNA (At3g41768) genes.

### Analysis of the biogenesis of organellar respiratory chain complexes in *msp1* mutants

The respiratory machinery in plants is composed of four major electron transport carriers, denoted as complexes I to IV (CI-CIV), the ATP synthase (CV), and several enzymes involved in non-phosphorylating bypasses of the electron transport chain (ETC), as alternative oxidases (AOXs) and rotenone-insensitive NAD(P)H dehydrogenases (NDs) (Jacoby, *et al*. 2012, Millar, *et al*. 2011, Schertl and Braun 2014, Senkler *et al*. 2017, Subrahmanian *et al*. 2016). Various studies indicate that altered mtRNA metabolism in plants may have strong effects on mitochondria biogenesis and function (see e.g., (Brown, *et al*. 2014, Colas des Francs-Small and Small 2014, Zmudjak and Ostersetzer-Biran 2017). The strong defects we observed in the processing *nad1* pre-RNAs (Figs. 5 and 6) suggest that NAD1 exists in extremely low levels or is completely absent from the mitochondria of *msp1-1* mutant-plants. The relative accumulation (*i.e*. steady-state levels) of native respiratory complexes, in 3-week-old wild-type (Col-0) plants, *nmat1-1* mutants (Keren, *et al*. 2012), rescued *msp1-1* seedlings, and Col-0 plantlets obtained from immature seeds (*i.e.*, at the torpedo stage), were analyzed by Blue-native (BN) PAGE analyses, followed by immunoblots assays (see Materials and Methods). These analyses indicated that the respiratory chain complex I (CI) is below detectable levels in *msp1-1* mutant (Fig. 7, upper panel). In-gel activity assays further showed that no complex I activity could be detected in the rescued *msp1-1* plants, while the accumulation and activity of CI was not affected in plantlets obtained from immature seeds of wild-type collected at the torpedo stage.

**Figure 7.**
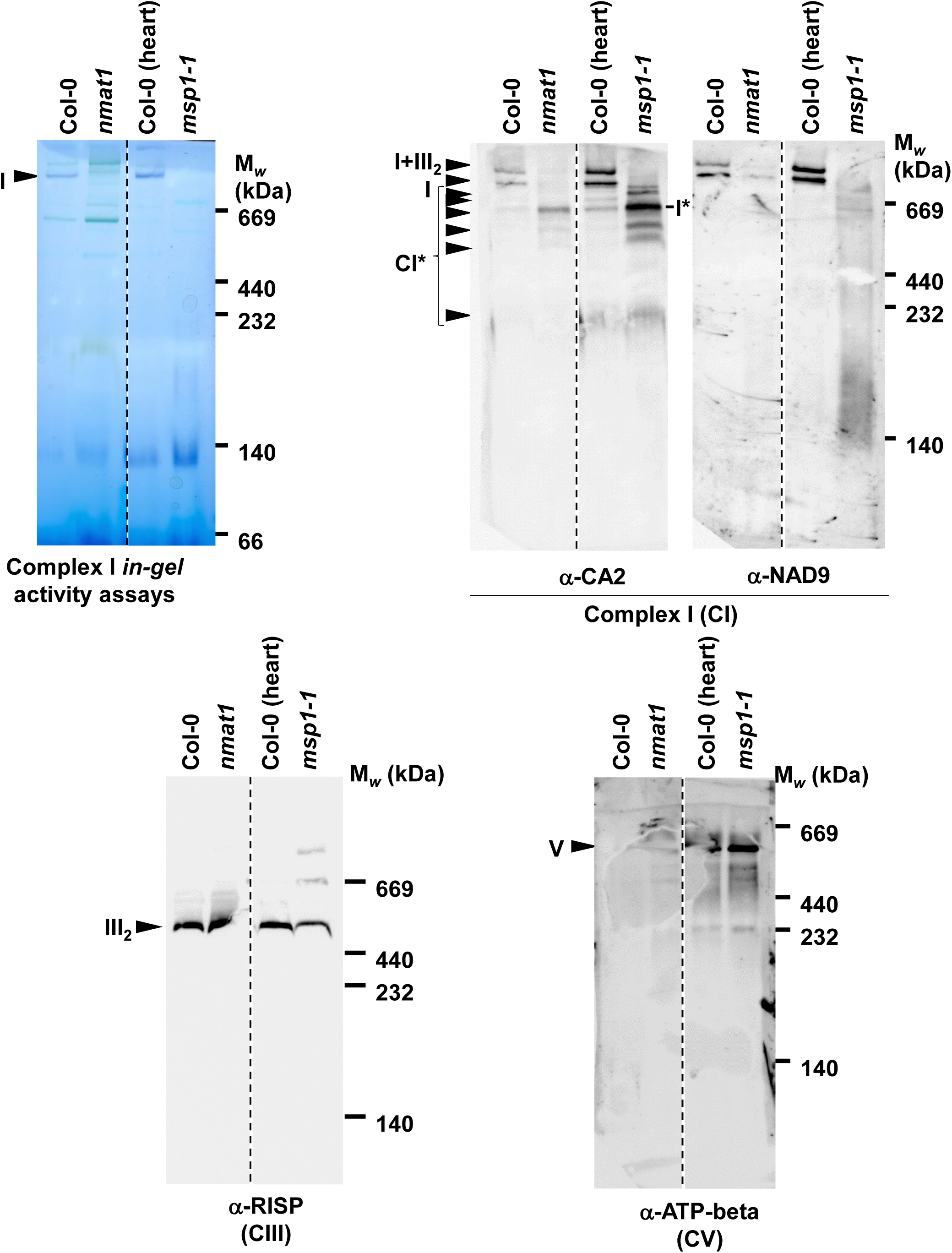
Relative accumulation of organellar proteins in wild-type plants and *msp1* mutants. Blue-native (BN) PAGE analyses. Blue native (BN)-PAGE of crude organellar membranous fractions was performed generally according to the method described by Pineau, *et al*. (2008). An aliquot equivalent to 40 mg of crude Arabidopsis mitochondria extracts, obtained from wild-type and *msp1* plants was solubilized with 5% (w/v) digitonin, and the organellar complexes were resolved by BN-PAGE. For immunodetection, the proteins were transferred from the native gels onto a PVDF membrane. The membranes were distained with ethanol before probing with specific antibodies (Table S3), as indicated below each blot. In-gel complex I activity assays were performed essentially as described previously (Meyer, *et al*. 2009). Arrows indicate to the native supercomplex I+III2 (∼1,500 kDa), holo-complexes I (∼1,000 kDa), III2 (∼500 kDa) and V (∼600 kDa). The asterisk in the CA2 panel indicates to the presence of various bands (*i.e*., labeled as C*, with calculated masses of about 800, 750, 700, 650, 600 and 250 kDa, respectively) which corresponds to a complex I assembly intermediates. Note that the four lanes in each of the immunoblots were loaded and separated on the same gels. Yet, we cropped to images to remove lanes corresponding to another PPR mutant that is unrelated to *msp1*. No other modifications were done to the original images.

The immunoblots with antibodies to CA2 subunit of the membranous arm of CI further indicated the accumulation of lower molecular weight bands (*i.e*., labeled as C*, with calculated masses of about 800, 750, 700, 650, 600 and 250 kDa, respectively) in *msp1-1* mitochondria, which most likely correspond to complex I assembly intermediates (Braun *et al*. 2014, Meyer 2012, Meyer *et al*. 2011). A band of a molecular mass of ∼700 kDa (labeled as I*) was also apparent in the immunoblots with antibodies raised to the soluble domain NAD9 subunit of complex I *nmat1* (Keren, *et al*. 2012), further supporting that the protein-bands seen in the CA2-blot correspond to sub-CI particles. No significant changes in the steady-state levels of complexes III and V were apparent in *msp1* mutant, as indicated by immunoblots with antibodies to RISP and ATP-B subunits, respectively. Arrows indicate to the native holo-complexes I (∼1,000 kDa), III (dimer, ∼500 kDa), V (∼600 kDa) and the supercomplex I+III_2_ (about 1,500 kDa). The protein profiles of *msp1* mutants correlate with several other Arabidopsis mutants that are lacking complex I activity, which are strongly affected in their cellular physiology but are able to germinate *in vitro* on sugar-containing media supplemented with various cofactors and nutrients (Dahan, *et al*. 2014, Fromm, *et al*. 2016a, Kuhn, *et al*. 2015, Ostersetzer-Biran 2016).

### Analyses of the mitochondrial respiratory activity of *msp1* plants

To analyze whether the respiratory activity was altered in *msp1* mutant, the oxygen-uptake rates of 200 mg leaf-disks obtained from 3-week-old MS-grown wild-type plants and *nmat1* (Keren, *et al*. 2012, Nakagawa and Sakurai 2006), embryo rescued homozygous *msp1-1* plantlets (*msp1* mutant lines) and heterozygous SALK-142675 plants, were monitored in the dark with a Clark-type electrode (Fig. 8). The average O_2_-uptake rates of *nmat1-1* mutants (Keren, *et al*. 2012) (109.81 ± 10.68 nmol O_2_ min^−1^ grFW^−1^) and rescued *msp1-1* plantlets (65.22 ± 11.34 nmol O_2_ min^−1^ grFW^−1^) were notably lower (i.e., about x2 to x3) than those of Col-0 (176.48 ± 12.36 nmol O_2_ min^−1^ grFW^−1^) or heterozygous *msp1* line (175.93 ± 26.23 nmol O_2_ min^−1^ grFW^−1^) (Fig. 8, Mock). Our own data and published results show that Arabidopsis mutants with no, or very low, levels of complex I often undergo oxidative stress and show a strong upregulation in the expression (and accumulation) of various NDs and AOXs proteins (Fromm, *et al*. 2016a, Karpova *et al*. 2002, Keren, *et al*. 2012, Kuhn, *et al*. 2015, Ostersetzer-Biran 2016). Accordingly, the relative accumulations of various AOXs and NDs, in particularly AOX1a and NDB2, are notably higher in *msp1* mutants than in the wild-type plants (Fig. 6b).

**Figure 8.**
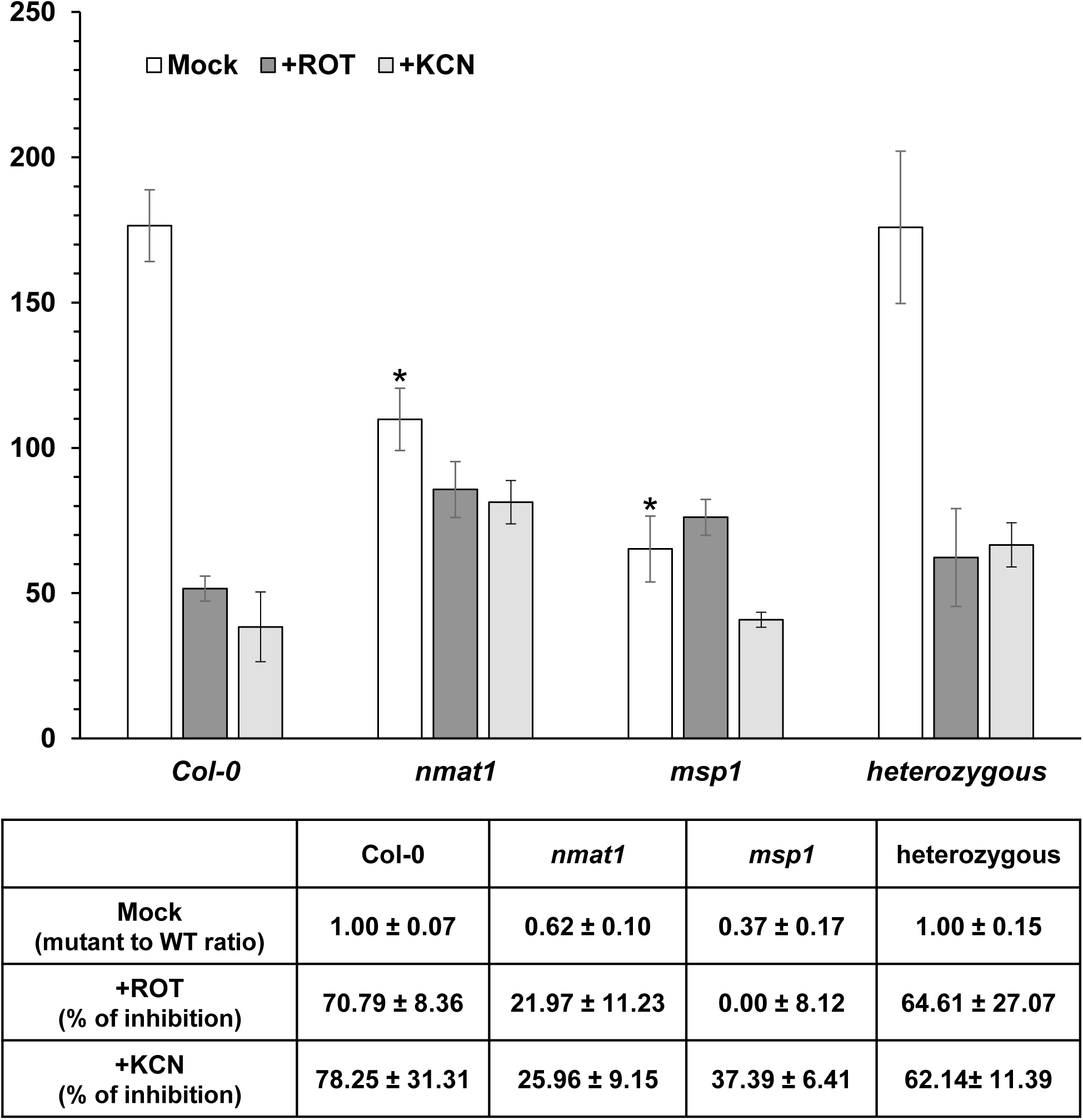
Respiration activities in wild-type and mutant lines. O_2_-uptake rates of wild-type plants and *nmat1* (Keren, *et al*. 2012, Nakagawa and Sakurai 2006), embryo rescued homozygous *msp1-1* plantlets (*msp1* mutant lines) and heterozygous SALK-142675 plants, were analyzed with a Clark-type electrode as described previously (Cohen, *et al*. 2014, Zmudjak, *et al*. 2017). For each assay, 200 mg MS-grown seedlings were submerged in 2.0 mL sterilized water and applied to the electrode in a sealed glass chamber in the dark. O_2_-uptake rates were measured in the absence (Control) or in presence of rotenone (+ROT, 50 μM) and KCN (1 mM) which inhibit complexes I and IV activities (respectively). The values are means of three to four biological replicates (error bars indicate one standard deviation). Asterisks indicate a significant difference from wild-type plants (Student’s T-test, P 0.05).

Pre-incubation of the plants with 50 μM rotenone (Maliandi *et al*. 2015), strongly affected the respiration rates of wild-type and the heterozygous *msp1* plants (*i.e*. 52.32 ± 7.25 and 62.26 ± nmol O_2_ min^−1^ gr FW^−1^, respectively), about 65 to 70% lower than the O_2_-upatek rates under the standard conditions (Fig. 8, +ROT). However, rotenone appeared to have much less effect on the respiratory activity of *nmat1* plants (*i.e*. 85.68 ± 9.62 nmol O_2_ min^−1^ grFW^−1^), while no obvious effect on the respiration activity of homozygous *msp1* plants were seen following the application of the inhibitor to the plants (*i.e.*, 76.12 ± 6.18 nmol O_2_ min^−1^ grFW^−1^) (Fig 8, +ROT). We also measured the O_2_-uptake rates of wild-type and mutant seedlings in the presence of potassium cyanide (KCN, 1 mM), which inhibits electrons transport through complex IV. The data shown in Figure 8 indicate that the mitochondrial electron transport inhibition by CN^−^ is much less pronounced in *nmat1* (81.30 ± 7.44 nmol O_2_ min^−1^ grFW^−1^, 26.0% inhibition) or *msp1* mutants (40.84 ± 2.62 nmol O_2_ min^−1^ grFW^−1^, 37.4% inhibition), whereas the O_2_-uptake rates of wild-type (38.38 ± 12.02 nmol O_2_ min^−1^ grFW^−1^) and heterozygous *msp1* plants (66.60 ± 7.59 nmol O_2_ min^−1^ grFW^−1^) were strongly affected in the presence of CN^−^ (78.3% and 62.1% inhibition, respectively). These data suggested that CI biogenesis and activity are strongly affected in the homozygous *nmat1* and *msp1* mutant-lines. Reduced sensitivities to rotenone and cyanide are tightly associated with an upregulation in the expression of proteins of the alternative electron transport system, including rotenone-insensitive NDs and CN-resistant AOXs (Fig. 6a), which can bypass complexes I and IV, respectively

## Discussion

### At4g20090 gene-locus encodes an essential PPR protein that is pivotal for the maturation of *nad1* pre-RNA transcripts in Arabidopsis mitochondria

Proteins that facilitate the splicing of the organellar group II introns are classified into two main categories, based on their topologies and predicted evolutionary origins (Barkan 2004, Brown, *et al*. 2014, Lambowitz *et al*. 1999, Schmitz-Linneweber, *et al*. 2015). These include maturases, which are encoded within the introns themselves and facilitate the splicing of their own cognate intron-RNAs, and various *trans*-acting cofactors, many of which cannot be found in the proposed bacterial progenitors of the mitochondria, and were thus likely recruited from the hosts during the endosymbiosis event to function in the processing of organellar group II introns (Brown, *et al*. 2014, Schmitz-Linneweber, *et al*. 2015, Zmudjak and Ostersetzer-Biran 2017).

In this work, we provide with genetic and biochemical evidences that a P-subclass PPR protein, EMB1025 encoded by the At4g20090 gene-locus, is essential for the processing of the *trans*-spliced *nad1* intron 1 in Arabidopsis mitochondria. *EMB1025* is found in the Arabidopsis SeedGenes database (http://seedgenes.org/) among other genes that are postulated to have essential functions during embryo development. Macroscopic analyses of siliques obtained from a heterozygous *msp1-1* line (SALK-142675) indicated a premature arrest of embryo development at the torpedo stage (Fig. 3). Here, we changed the annotation of EMB1025 into MSP1 (Mitochondria Splicing PPR-factor 1) to better indicate the established molecular functions of At4g20090 in group II intron RNA splicing in Arabidopsis mitochondria. *In silico* prediction databases suggest that *MSP1* encodes a mitochondria-localized protein, which harbors 15 repeats of the degenerated PPR-motif (Figs. 1, S1, S2 and Table S1). In transformant tobacco cells, the signal of the MSP1-GFP fusion protein is localized to mitochondria (Fig. 2 and (Lurin, *et al*. 2004).

PPR proteins are pivotal in the posttranscriptional regulation of gene-expression in plant mitochondria (reviewed in *e.g*., (Barkan and Small 2014, Colas des Francs-Small and Small 2014, Dahan and Mireau 2013, Gorchs Rovira and Smith 2019, Hammani and Giege 2014, Zmudjak and Ostersetzer-Biran 2017). Many PPRs show gene-specific expression patterns, during plant growth and development and various stresses stimuli (Chen, *et al*. 2018, Lee and Kang 2016, Xing *et al*. 2018). These may therefore serve as fundamental players in the complex molecular crosstalk between the nucleus and the mitochondria, linking organellar functions with developmental and/or environmental signals. Various *ppr* mutants where previously shown to affect the splicing of mitochondrial group II introns found in either *cis-* or *trans*-configurations (Colas des Francs-Small, *et al*. 2014, Colas des Francs-Small and Small 2014, Falcon de Longevialle, *et al*. 2007, Gualberto, *et al*. 2015, Hammani and Giege 2014, Koprivova, *et al*. 2010, Liu, *et al*. 2010, Schmitz-Linneweber, *et al*. 2015, Wang, *et al*. 2017, Wang, *et al*. 2018, Weissenberger, *et al*. 2017, Yang, *et al*. 2014, Zmudjak and Ostersetzer-Biran 2017). We sought to define the molecular functions of MSP1 and to analyze its postulated roles in mitochondria RNA-metabolism, by establishing loss-of-function mutations in the At4g20090 gene-locus.

As no homozygous mutant individuals were identified among the mature seeds of self-fertilized heterozygous *msp1* plants, we developed an original embryo rescue approach (see Materials and Methods), modified from methods used to recover germination-arrested phenotypes in Arabidopsis plants (Cordoba, *et al*. 2016, Dahan, *et al*. 2014, Franzmann, *et al*. 1989, Fromm, *et al*. 2016a). This strategy allowed us to obtain homozygous *msp1-1* plantlets *in vitro*, which displayed low germination and severe developmental defect phenotypes (Fig. 4). Analyses of the mtRNA patterns in wild-type and rescued homozygous *msp1* seedlings indicated a strong perturbation in the maturation of *nad1* intron 1 pre-RNAs, where the levels of *nad1* mRNA was found to be reduced by about 10,000 folds (Fig. 5). The data also indicated to reduce steady-state levels of mature transcripts corresponding to *nad2* exons a-b and exons c-d, and *rpl2* mRNA.

The genes of *nad1*, *nad2* and *rpl2* are all interrupted by group II intron sequences in the mtDNA of Arabidopsis. We thus analyzed whether the reduced levels of *nad1*, *nad2* and *rpl2* mRNA in the mutants may relate to splicing defects in the absence of a functional MSP1 protein. When we compared the ratios of pre-RNAs to mRNAs of the 23 introns in Arabidopsis mitochondria, between *msp1* and wild-type plants, a strong defect in the maturation of *nad1* intron 1 was apparent in the mutant (*i.e.*, ∼250,000 folds) (Fig. 6a). A reduced splicing efficiency was also seen for *nad2* i1 (*i.e*., about 4x lower) (Fig. 6a). Yet, given the small extent to which this mRNA was reduced in the mutant (Fig. 5a), and the fact that the splicing of various *nad2* introns are also interrupted in other Arabidopsis mutants affected in mtRNA metabolism (Falcon de Longevialle, *et al*. 2007, Keren, *et al*. 2009), it is difficult to draw firm conclusions about the roles of MSP1 in the splicing of this intron. As the splicing efficiencies of *nad2* i3 and *rpl2* i1 were found similar in *msp1-1* mutant and the wild-type plantlets (Fig. 6a), we concluded that the splicing of these introns does not rely upon MSP1. No significant changes in the steady-state levels of various intronless transcripts (Fig. 5) or the splicing efficiencies in the remaining 19 introns were observed in the mitochondria of *msp1-1* (Fig. 6a). Taken together, the RNA profiles in wild-type and *msp1-1* mutants (Figs. 5 and 6) indicate a key role for MSP1 in the *trans*-splicing of *nad1* intron1 in Arabidopsis mitochondria. The RNA profiles seem specific to the functions of MSP1, as wild-type plantlets obtained from immature seeds at the same embryo developmental (*i.e*., torpedo) stage, which grown together with *msp1-1* line, did not show any obvious defects in the accumulation or processing of various other organellar transcripts (Fig. 5b).

### Impaired complex I biogenesis in *msp1* mutants

The steady-state levels of the proteins is accomplished largely by the balance between the rate of translation and protein degradation. Defects in mtRNA metabolism are expected to result with altered translation efficiencies. Complexes, I, III, IV, V and the ribosomes are all assemblies of subunits that are expressed by both nuclear-and organellar loci, thus necessitating complex mechanism to regulate the expression and accumulation of the organellar-localized proteins during development and various growth conditions (Gualberto *et al*. 2014, Hammani and Giege 2014, Law *et al*. 2014, Zmudjak and Ostersetzer-Biran 2017). Accordingly, organellar ribosome footprint analyses indicate that mitochondria-encoded mRNAs are differentially translated, while the synthesis of plastidial proteins directly correlate with the abundances of the mature transcript with the organelles (Chotewutmontri and Barkan 2016, Planchard, *et al*. 2018).

NAD1 is a central anchor component of mitochondrial complex I (Badri *et al*. 2013, Baradaran *et al*. 2013, Braun, *et al*. 2014, Fromm *et al*. 2016b, Klodmann *et al*. 2010, Kuhn, *et al*. 2015, Lazarou *et al*. 2009, Meyer 2012, Shimada *et al*. 2014, Subrahmanian, *et al*. 2016). A strong reduction in NAD1 is known to affect the assembly and stability of CI, resulting with altered plant growth and development (Cohen, *et al*. 2014, Falcon de Longevialle, *et al*. 2007, Keren, *et al*. 2012, Ren *et al*. 2017, Wang, *et al*. 2017). Analysis of the protein profiles indicated that holo-complex I and the supercomplex I+III_2_ are below detectable levels in *msp1* plants (Fig. 7), whereas the accumulation of other respiratory complexes (*i.e*., native complexes III and V) was not significantly affected in the mutant (Fig. 7). In-gel activity and BN-PAGE/immunoblot assays indicated that CI is significantly reduced yet still visible in *nmat1* mutant (Fig. 7), which is also affected in the splicing of *nad1* i1, but seem to accumulates *nad1* mRNA to higher levels than *msp1* line (Keren, *et al*. 2012). We further noticed the accumulation of lower molecular weight bands in *msp1-1* mitochondria (Fig. 7, labeled as C*), which are likely associated with partially assembled CI particles (Braun, *et al*. 2014, Cohen, *et al*. 2014, Keren, *et al*. 2012, Meyer 2012, Meyer, *et al*. 2011). The 700 kDa band, which is also apparent in *nmat1* mutant, was found to accumulate to high levels in *msp1* and seems also apparent in the immunoblots with NAD9 protein (Fig. 7, labeled as I*). Thus, differently from previous reports suggesting that NAD1 is required early in CI assembly (*i.e*., during the formation of the 450 subdomain of the membrane-arm) (Meyer 2012, Meyer, *et al*. 2011, Subrahmanian, *et al*. 2016), these data may indicate that NAD1 rather incorporates into CI, into an assembly intermediate which already contains at least some subunits of the soluble arm (Fig. 7, NAD9 panel).

To examine the effects of CI loss, or its reduction to trace amounts, on the respiratory activities of *msp1* mutants, we measured the O_2_-uptake rates of Arabidopsis wild-type and mutant plants by a Clark-type electrode. Notable reductions in the O_2_-uptake rates were observed between the *nmat1* (38% lower) and *msp1* (63% lower) mutant-lines and the wild type plants (Fig. 8). Inhibition of mitochondrial respiratory chain complex I by rotenone had a stronger effect on wild-type and heterozygous *msp1-1* plants (65-70% reduced activities, respectively) than on *nmat1* (20% reduction) and *msp1-1* mutant (no reduction in the O_2_-uptake rates) (Fig. 8), which are both affected in CI biogenesis (Figure 7).

We further analyzed the respiratory activities in the presence of KCN, a complex IV-specific inhibitor. As shown in Figure 8, the presence of cyanide also had a stronger effect on wild-type plants (about 80% decreased O_2_-uptake rates) and heterozygous *msp1-1* plants (about 60% lower respiration activity), whereas the respiration rates were decreased by only 26% and 37% in *nmat1* and *msp1* (respectively) in the presence of the inhibitor. These results are correlated with several mutants affected in RNA metabolism and the expression of complex I subunits (*e.g*., (Cohen, *et al*. 2014, Dahan, *et al*. 2014, Keren, *et al*. 2012), but are different from those of *ndufv1* mutants which rather exhibit increased O_2_-uptake rates (Kuhn, *et al*. 2015). The increased respiration in *ndufv1* is associated with the induction of alternative pathways of electron transport, via CN*-* resistant alternative oxidases (AOXs) and rotenone insensitive type-II NAD(P)H dehydrogenases (NDs) (Araújo *et al*. 2012, Jacoby, *et al*. 2012, Juszczuk *et al*. 2012, Millar, *et al*. 2011, Rasmusson *et al*. 2008, Van Aken *et al*. 2009, Yoshida *et al*. 2008). RT-qPCR analyses indicated that the relative steady-state levels of various AOXs and NDs, in particularly AOX1a and NDB2, are notably higher in *msp1* mutants than in the wild-type plants (Fig. 6b). Therefore, the molecular basis for the differences in the respiration activities between *msp1* and *ndufv1* mutants remains unclear, as both plant lines seem to lack CI (or possessing only trace amounts of the native complex) and accumulate high levels of AOXs and NDs. It remains to be addressed why *msp1* mutants seems more severely affected in development, and to whether these lines show different patterns in the regulation of respiratory fluxes and metabolic switches (*i.e*., the tricarboxylic acid, glycolysis, lipids or amino acid metabolism).

### Morphology of *MSP1* mutant alleles

Plant embryogenesis is a highly complex developmental process. Some Arabidopsis mutants with impaired embryogenesis (*emb*) at the heart to cotyledon stages of development, can be sometimes rescued on a nutrient medium designed to promote plant regeneration from immature wild-type cotyledons (Franzmann, *et al*. 1989). Many of the rescued mutants, exhibit abnormal vegetative and reproductive development. Mutants arrested at earlier stages in embryogenesis (*e.g*., globular or triangular stages) can be also rescued, but these often produce callus, but fail to develop into mature seedlings (Franzmann, *et al*. 1989). Mitochondria have pivotal roles in cellular metabolism, even in anaerobic organisms which relay, for at least part of their life cycle on glycolysis for ATP synthesis (Muller *et al*. 2012). Altered mitochondrial functions in plants are commonly associated with low germination and various growth to developmental defect phenotypes. Previously, it has been noted that complex I defects in plants can result in a broad spectrum of phenotypes, ranging from mild growth to severe developmental defects and arrested embryogenesis (Dahan, *et al*. 2014, Kuhn, *et al*. 2015, Ostersetzer-Biran 2016). In some rear cases plants may adapted to live in the absence of complex I, as the parasitic mistletoe (*Viscum album*) plant (Maclean *et al*. 2018, Petersen *et al*. 2015, Senkler *et al*. 2018).

Arabidopsis mutants affected in complex I biogenesis and function are generally able to produce viable homozygote mutants, while the functions of complexes III, IV and the translation machinery are regarded to be essential. Processes that seem particularly critical to the establishment of Arabidopsis mitochondrial mutants include germination, root development and flowering. Embryo development, and probably the ability to penetrate through the seed coat (Radchuk and Borisjuk 2014), requires high amounts of metabolic energy, which is provided during seed-development through the degradation of reserves stored in the seed-endosperm. In some plants, including *Arabidopsis thaliana*, the cotyledons absorb much of the nutrients from the endosperm and thus serve as the main energy supply source for the embryo. Accordingly, embryo size in Arabidopsis is strongly influenced by the size of the endosperm prior to its breakdown and absorbance by the cotyledons (Fourquin *et al*. 2016). Accordingly, severe complex I mutants, as *nma1*, *nmat4*, *ndufv1* (Cohen, *et al*. 2014, Keren, *et al*. 2012, Kuhn, *et al*. 2015, Nakagawa and Sakurai 2006) and *msp1* (Fig. 3), are strongly impaired in the storage of sugar and oil reserve. Analyses of immature siliques in their heterozygous lines indicate to about 25% wrinkled brown seeds. Genotyping indicate that these individuals are homozygous for the mutations. Accordingly, examination of the seed morphology of heterozygous *msp1* plants, indicated to about 25% white-translucent seeds which later developed into shrunken/wrinkled brown seeds (see Fig. 3). Interestingly, their structures resemble that of their arrested embryo-developmental stage (*i.e.*, torpedo or ‘walking-stick’ stages of embryogenesis) (Fig. 3a).

Altered energy reserves are likely affecting seed quality, which relate to the variability observed between individuals of the same progenies in severe mutant lines (Cohen, *et al*. 2014, Kuhn, *et al*. 2015) (Figs. 3 and 4). When the seedlings were transferred from the MS-media to the soil, they develop small rosettes (Fig. 4f) with curled leaves, a leaf-defect phenotype that is also noted in other Arabidopsis mutants affected in mitochondria biogenesis (Córdoba *et al*. 2019, Falcon de Longevialle, *et al*. 2007, Gibala *et al*. 2009, Hsieh *et al*. 2015, Radin *et al*. 2015). While root development is compromised in several mutants affected in mitochondria biogenesis (Cohen, *et al*. 2014, Dahan, *et al*. 2014, Hsieh, *et al*. 2015, Keren, *et al*. 2012, Nakagawa and Sakurai 2006), the *msp1* mutants rather developed a normal root system (Fig. 4e). These differences may relate to the media used to rescue the *msp1* mutants (see Material and Methods).

Why does the loss of *msp1* cause a major embryo defect, while the seeds of homozygous *nmat1*, *nmat4* and *otp43* mutants, which are also strongly affected the processing of *nad1* intron 1 are viable and able to germinate on soil? Currently, we cannot provide a definitive explanation, but we speculate that such variations in growth and developmental defect phenotypes, observed between these mutants, may relate to the presence of residual level of NAD1 protein in *nmat1*, *nmat4* and *otp43* mutants (*i.e*., even a small accumulation of NAD1 protein may have a strong effect during embryogenesis and at later developmental stages). It remains possible that the severity of the growth phenotypes is directly related with the levels of NAD1. Mutants with no complex I activity may fail to germinate due to insufficient energy supply, or might be unable to penetrate the hard seed-coat found in the mature seeds (Kuhn, *et al*. 2015).

Alternatively, it remains possible that MSP1 may also affect RNA processing events other than the splicing of *nad1* i1. Such ‘off-targets’ may relate to the processing of additional mitochondrial tRNAs, rRNAs or non-coding RNAs, which were not included in the transcriptome and splicing assays (Figs. 5 and 6). Canonical PPR proteins are sequence-specific RNA-binding factors, which affect the processing of a single, or a few, closely related transcripts (Barkan, *et al*. 2012, Barkan and Small 2014, Coquille, *et al*. 2014, Takenaka *et al*. 2013, Yagi, *et al*. 2013, Yagi, *et al*. 2014). A combinatorial code for RNA-recognition by PPR proteins was recently suggested, based on amino acids found at positions 5 and 35 that are postulated to be specificity-determining positions for the association with RNA (Barkan, *et al*. 2012, Cheng *et al*. 2016, Takenaka, *et al*. 2013). According to the PPR-binding code (PPRCODE server; (Shen *et al*. 2019), the 15 suggested PPR motifs in MSP1 have a predicted RNA-binding sequence of 5’-N-N-(C>U)-(U>C>G)-A-(C>U)-(C>U)-(A>C)-A-A-A-(G>>C)-A-(U>C>G)-N-3’ (N indicates unknown bases) (Suppl. Fig. S1, Table S1). A sequence region with a high homology to the predicted MSP1-binding site (i.e., 5’-AG**<u>UCAU</u>**A**<u>AAAAGAU</u>**A, nucleotides 82701-82715) was identified in BLAST search, using the up-to-date RefSeq of Arabidopsis mtDNA (accession BK010421) (Sloan *et al*. 2018). Interestingly, this sequence corresponds to *nad1* intron 1, the genetically-identified intron target of MSP1 protein, but no other regions with significant homology to the predicted MSP1-binding site could be identified in the mtDNA sequence of *Arabidopsis thaliana* Col-O.

The structure of *nad1 i1* was modeled based on a previous structural models of group II intron RNAs (Candales *et al*. 2011, Dai *et al*. 2003), structural analyses of bacterial MATs bound to their cognate group II intron RNAs (Piccirilli and Staley 2016, Qu, *et al*. 2016, Zhao and Pyle 2016), and the secondary structure prediction algorithms Mfold (Zuker 2003) and Alifold (Hofacker *et al*. 2002). Figure 9 highlights the predicted binding site of MSP1 on the secondary structure model of nad1 intron 1 RNA sequence. Nucleotides predicted to be involved in the canonical group II introns tertiary interactions, including the IBS1-EBS1, IBS2-EBS2, α-α’, β-β’, γ-γ’, δ-δ’, ε-ε’, ζ-ζ’, κ-κ’, λ-λ’, θ-θ’ and the η-η’ sites, are indicated in Figure 9.

**Figure 9.**
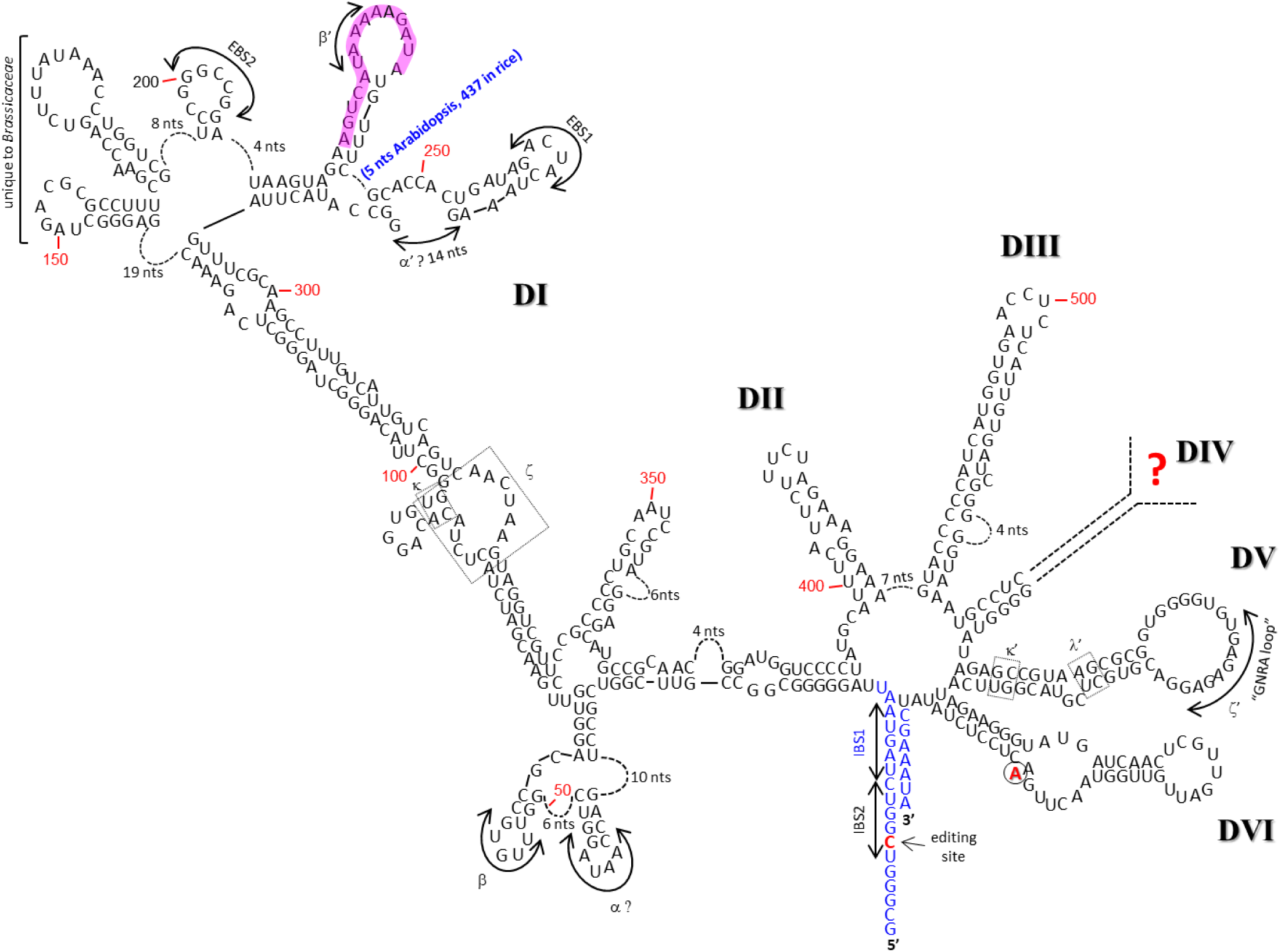
Secondary structure model of the Arabidopsis *nad1* intron 1 with the predicted MSP1 protein binding sequence region. The sequence of *Arabidopsis thaliana* (Col-0) *nad1* intron 1 includes 16 nucleotides of the 5’ exon, 11 nucleotides of the 3’ exon, 481 nucleotides of the *trans*-encoded 5’-part of the intron and 108 nucleotides of its *trans*-encoded 3’ part, with residue numbering beginning at the first intron nucleotide (*i.e.*, 5’-part). The structure of the intron was modeled based on a previous structural models of group II intron RNAs (Candales, *et al*. 2011, Dai, *et al*. 2003), structural analyses of bacterial MATs bound to their cognate group II intron RNAs (Piccirilli and Staley 2016, Qu, *et al*. 2016, Zhao and Pyle 2016), and the secondary structure prediction algorithms MFOLD (Zuker 2003) and Alifold (Hofacker, *et al*. 2002). Nucleotides predicted to be involved in the canonical group II introns tertiary interactions, including the IBS1-EBS1, IBS2-EBS2, α-α’, β-β’, γ-γ’, δ-δ’, ε-ε’, ζ-ζ’, κ-κ’, λ-λ’, θ-θ’ and the η-η’ sites are indicated. The predicated cleavage site, with the bulged A, is indicated according to (Li-Pook-Than and Bonen 2006). The predicted MSP1 binding site (*i.e*., 5’-AGUCAUAAAAAGAUA) is highlighted (magenta color) on the predicated 2D structure of the Arabidopsis *nad1* intron 1.

According to the predicted RNA-binding site and the putative 2D structure of *nad1* intron1, it is postulated that MSP1 binds to a stem-loop region corresponding to the β’ site (Fig. 9, highlighted in magenta color), a region that is highly important for the folding of group II introns into their catalytically active form. It is expected that MSP1, as other PPR proteins, would bind to a short single-stranded loop region. The association of MSP1 with this loop region might promote tertiary interactions within the intron that are important for RNA catalysis, as indicated in the case of the group II intron-encoded maturase proteins (Piccirilli and Staley 2016, Qu, *et al*. 2016, Zhao and Pyle 2016).

## Materials and Methods

### Plant material and growth conditions

*Arabidopsis thaliana* (ecotype Columbia) was used in all experiments. Wild-type (Col-0) and *mps1* mutant lines SALK-142675 (*msp1-1*), SALK-018927 (*msp1-2*) and SALK-070654 (*msp1-3*) were obtained from the Arabidopsis Biological Resource Center (ABRC) at Ohio State University (Columbus, OH). Prior to germination, seeds obtained from wild-type and mutant lines were surface-sterilized with Cl_2_ gas, generated by the addition of 1 ml HCl per 30 mL of bleach (sodium hypochlorite 4.7%), for 4 hours at room-temperature (RT). The seeds were than sown on MS-agar plates containing 1% (w/v) sucrose or rescued by a method described in detail below. For well-synchronized germinations, the seeds were kept in the dark for 5 days at 4°C and then grown under long day condition (LD, 16:8-hour) in a controlled temperature and light growth chamber (Percival Scientific, Perry, IA, USA) at 22°C and light intensity of 300 µE m^−2^ s^−1^. PCR was used to screen the plant collection and check the insertion integrity of each individual line (specific oligonucleotides are listed in Suppl. Table S2). Sequencing of specific PCR products was used to analyze the precise insertion site in the T-DNA lines.

### GFP localization assays

In-vivo localizations by GFP-fusion protein assays were performed essentially as described previously (Shevtsov *et al*. 2018b). A 450 nucleotide fragment of the 5’-terminus of MSP1 (At4g20090) was fused in-frame to eGFP, and cloned into pSAT6 vector (Tzfira *et al*. 2005). The resulting construct (*N’*-MSP1-GFP) was cloned into the binary pCAMBIA-2300 vector (Specific primers are listed in Suppl. Table S2). The vector was introduced into *Agrobacterium tumefaciens* (strain EHA105), and the cells were grown overnight at 28°C in YEP medium supplemented with kanamycin (50 mg/L). The cells were harvested by centrifugation, resuspended to a final concentration equivalent to A_600_ ∼ 1.0 O.D. in transformation buffer (Shevtsov, *et al*. 2018b). For localization analyses, 3∼5 young leaves of *N. benthamiana* (4 to 6 week-old) were infiltrated with the Agrobacterium suspension using a 1 mL syringe (abaxial side, without a needle). Constructs containing the GFP protein alone was used as a control. After 24∼48 hours, the Agro-infiltrated leaves were analyzed by confocal microscopy (Olympus IX 81, Fluoview 500, Imaging facility, Volcani center). To visualize mitochondria in vivo, plant tissue was incubated with 2 µM MitoTracker^®^ for 10 min at room temperature prior to the confocal analyses.

### Embryo rescue and establishment of homozygous *msp1* mutants

Siliques, obtained from wild-type and heterozygous SALK-142675 (*msp1-1*) plants, were surfaced sterilized with 6% bleach solution for 10 min at RT. The seeds were then soaked in a 70% ethanol solution for 10 min at RT, washed briefly with sterile DDW, and opened in a biological hood. Green (*i.e.*, wild-type or heterozygous) and white *msp1-1* mutant seeds (homozygous ones), were obtained from siliques of heterozygous SALK-142675 (*msp1-1*), at different developmental stages (i.e., distributed along the Inflorescence Stem). Likewise, immature seeds of Col-0 seeds at the torpedo stage were obtained from sterilized siliques. The seeds were then sowed on MS-agar plates supplemented with 1% (w/v) sucrose and different vitamins (*i.e*., 10 mg Myoinositol 100 μg Thiamine, 100 μg Pyridoxine, 100 μg nicotinic acid). For DNA and RNA analysis we used Arabidopsis wild-type and *msp1* plantlets at stage R6 (*i.e.*, 6 to 8 leaves) (Boyes, *et al*. 2001). To obtain larger quantities of plant material, plantlets at stage R6 grown on MS-agar plates, where transplanted into a MS-based liquid medium supplemented with 1 to 3% (w/v) sucrose and various vitamins (10 mg Myoinositol 100 μg Thiamine, 100 μg Pyridoxine, 100 μg nicotinic acid) and incubated in Arabidopsis growth chambers (Percival Scientific, Perry, IA, USA), under standard growth conditions with moderate (50∼100 RPM) shaking (i.e., long day conditions, at 22°C and light intensity of 300 µE m^−2^ s^−1^).

### Microscopic analyses of Arabidopsis wild-type and mutant plants

Analysis of whole plant morphology, roots, leaves, siliques and seeds of wild-type and mutant lines were examined under Stereoscopic (dissecting) microscope or light microscope at the Bio-Imaging unit of the Institute of Life Sciences (The Hebrew University of Jerusalem).

### RNA extraction and analysis

RNA extraction and analysis was performed essentially as described previously (Cohen, *et al*. 2014, Keren *et al*. 2011, Shevtsov, *et al*. 2018b, Sultan, *et al*. 2016, Zmudjak, *et al*. 2013). In brief, RNA was prepared from 50 mg seedlings grown on MS-agar plates supplemented with 1% sucrose, following standard TRIzol Reagent protocols (Ambion, Thermo Fisher Scientific) with additional phenol/chloroform extraction procedure. The RNA was treated with DNase I (RNase-free) (Ambion, Thermo Fisher Scientific) prior to its use in the assays. RT-qPCR was performed with specific oligonucleotides designed to exon-exon (mRNAs) regions, corresponding to numerous mitochondrial genes, and intron-exon regions (pre-mRNAs) within each of the 23 group II introns in *Arabidopsis thaliana* (Suppl. Table S3). Reverse transcription was carried out with the Superscript III reverse transcriptase (Invitrogen), using 1 - 2 µg of total RNA and 250 ng of a mixture of random hexanucleotides (Promega) and incubated for 50 min at 50°C. Reactions were stopped by 15 min incubation at 70°C and the RT samples served directly for real-time PCR. Quantitative PCR (qPCR) reactions were run on a LightCycler 480 (Roche), using 2.5 μL of LightCycler 480 SYBR Green I Master mix and 2.5 μM forward and reverse primers in a final volume of 5 µL. Reactions were performed in triplicate in the following conditions: pre-heating at 95°C for 10 min, followed by 40 cycles of 10 sec at 95°C, 10 sec at 58°C and 10 sec at 72°C. The nucleus-encoded 18S rRNA (At3g41768) and the mitochondrial 26S ribosomal rRNA subunit (ArthMr001) were used as reference genes in the qPCR analyses.

### Crude mitochondria preparation from MS-grown Arabidopsis seedlings

Crude mitochondria extracts were prepared essentially as described previously (Pineau *et al*. 2008, Shevtsov *et al*. 2018a). For the preparation of organellar extracts, about 200 mg of liquid MS-grown plantlets were harvested and homogenized in 2 ml of 75 mM MOPS-KOH, pH 7.6, 0.6 M sucrose, 4 mM EDTA, 0.2% polyvinylpyrrolidone-40, 8 mM L-cysteine, 0.2% bovine serum albumin and protease inhibitor cocktail ‘complete Mini’ from Roche Diagnostics GmbH (Mannheim, Germany). The lysate was filtrated through one layer of miracloth and centrifuged at 1,300 ×g for 4 min at 4ᵒC (to remove cell debris). The supernatant was then centrifuged at 22,000 ×g for 10 min at 4ᵒC. The resultant pellet contains thylakoid and mitochondrial membranes was washed twice with 1ml of wash buffer 37.5 mM MOPS-KOH, 0.3 M sucrose and 2mM EDTA, pH 7.6. Protein concentration was determined by the Bradford method (BioRad) according to the manufacturer’s protocol, with bovine serum albumin (BSA) used as a calibrator.

### Blue native (BN) electrophoresis for isolation of native organellar complexes

Blue native (BN)-PAGE of crude organellar membranous fractions was performed generally according to the method described by Pineau, *et al*. (2008). An aliquot equivalent to 40 mg of crude Arabidopsis mitochondria extracts, obtained from wild-type and *msp1* plants was solubilized with 5% (w/v) digitonin in BN-solubilization buffer (30 mM HEPES, pH 7.4, 150 mM potassium acetate, 10% [v/v] glycerol), and then incubated on ice for 30 min. The samples were centrifuged 8 min at 20,000 xg to pellet any insoluble and Serva Blue G (0.2% [v/v]) was added to the supernatant. The samples were then loaded onto a native 4 to 16% linear gradient gel. For ‘non-denaturing-PAGE’ immunoblotting, the gel was transferred to a PVDF membrane (Bio-Rad) in Cathode buffer (50 mM Tricine and 15 mM Bis-Tris-HCl, pH 7.0) for 16 h at 4°C at constant current of 40 mA. The blots where then incubated with antibodies against various organellar proteins, and detection was carried out by chemiluminescence assay (various primary antibodies used in these assays are listed Suppl. Table S4) after incubation with an appropriate horseradish peroxidase (HRP)-conjugated secondary antibody. For in-gel complex I activity the gels were washed several times with DDW and the activity assays were performed essentially as described previously (Meyer *et al*. 2009). In-gel complex I activity assays were performed essentially as described previously (Meyer, *et al*. 2009).

### Respiration activity

Oxygen consumption (*i.e.* O_2_ uptake) measurements were performed with a Clarke-type oxygen electrode, and the data feed was collected by Oxygraph-Plus version 1.01 software (Hansatech Instruments), as described previously (Shevtsov, *et al*. 2018b). The electrode was calibrated with oxygen-saturated water and by the addition of excess sodium dithionite for complete depletion of the oxygen in water housed in the electrode chamber. Equal weights (200 mg) of wild-type and *msp1* seedlings were immersed in water and incubated in the dark for a period of 30 min. Total respiration was measured at 25°C in the dark following the addition of the seedlings to 2.5 mL of sterilized tap water in the presence or absence of rotenone (50 μM) and KCN (1 mM).

## Acknowledgments

This work was supported by grants to O.O.B from the ‘Israeli Science Foundation’ (ISF grant no. 741/15).

## Supplementary Materials

**Supplementary Figure S1. The topology of MSP1 protein.**

(a) The deduced amino acid sequence of MSP1 (At4g20090). The postulated regions, corresponding to the mitochondrial targeting sequence (36∼37 amino acid long) is underlined and highlighted in magenta). The predicted cleavage sites of the Mitochondrial Processing Peptidase (MPP, 36 aa) or Intermediate Cleavage Peptidase 55 (ICP55, 37 aa) are indicated by box. The 15 PPR motifs (highlighted in blue) were predicted by the SMART (Letunic, *et al*. 2012), the Conserved Domain Database (CDD) (Marchler-Bauer, *et al*. 2003) and PPRCODE (Shen, *et al*. 2019) servers. (b) To get more of an insight on MSP1’s mode of action, in particular of RNA recognition and binding, we performed an atomic model of the protein using the Phyre^2^ server (Kelley and Sternberg 2009). The model structure of MSP1 (*i.e*. ribbon and surface views) were generated by the PyMol software suite (DeLano and Lam 2005). “i” and “iv” represent the ribbon structure from different angles, while panels “ii” and “iii” represent the hypothetical surface of MSP1 protein. The color code is red for negative values, white for near zero values, and blue for positive values.

**Supplementary Figure S2. *MSP1* gene expression patterns at different tissues and during various developmental stages.**

The expression patterns of MSP1 were analyzed by publicly available microarray and high throughput sequencing databases, including The Arabidopsis Information Resource, TAIR; http://www.arabidopsis.org) and Genevestigator analysis toolbox (Hruz, *et al*. 2008, Zimmermann, *et al*. 2004).

**Figure S3. The nucleotide sequence of MSP1 gene and the precise locations of each of the T-DNA-insertional sites.**

The nucleotides sequence of *MSP1*, encoded by At4g20090 gene-locus. Underlined letters indicate to the 5’ and 3’ untranslated regions (UTRs), as indicated by the RACE analysis and TAIR database, while uppercased letters represent the open reading frame of mTERF22. The position of T-DNA insertions in *msp1-1* (SALK-142675), *msp1-2* (SALK-018927) and *msp1-3* (SALK-070654).

**Supplementary Table S1.** Prediction of MSP1 PPR motifs and analysis of their RNA-binding code.

**Supplementary Table S2.** Oligonucleotides used in screening of individual T-DNA insertion lines in Arabidopsis and cloning of the MSP1-GFP gene-fusion construct.

**Supplementary Table S3.** Lists of oligonucleotides used for the analysis of the splicing profiles of wild-type and mutant plants by RT-qPCR experiments.

**Supplementary Table S4.** List of antibodies used for the analysis of wild-type and *msp1* mutants.

## Author contributions statement

<u>Corinne Best</u>: Plant growth and analysis, embryo rescue and establishment of mutant lines, Biochemical analysis of gene expression, BN-PAGE analysis, RNA isolation, DNA sequencing, analysis of the transcriptome and splicing profiles of Arabidopsis mitochondria by RT-qPCR, respiration analyses. <u>Michal Zmudjak</u>: Embryo rescue and establishment of mutant lines, RNA isolation and analysis of transcriptome profiles of Arabidopsis mitochondria by RT-qPCR. <u>Prof. Oren Ostersetzer-Biran</u>: Manuscript preparation and Corresponding author.

