## Supplementary Data for "The PPR-related splicing cofactor MSP1/EMB1025 protein, encoded by At4g20090, encode an essential protein that is required for the splicing of *nad1* intron 1 and for the biogenesis of complex I in Arabidopsis mitochondria"

**a**

MPP/ICP55 cleavage sites

MPKCPIPIRISFFSYFLKESRILSSNPVNFSIHLRFSSSVSVSPNPSMEVVENPLEAPIS
EKMFKSAPKMGSFKLG|DSTLSSMIESYANSGDFDSVEKLLSRIRLENRVII|ERSFIVV
FRAYGKAHLPDKAVDLFHRMVDEFRCKR|SVKSFNSVLNVIINEGLYHRGLEFYDYVVNS
NMNMN|ISPNGLSFNLVIKALCKLRFVDRAIEVFRGMPERKCLPD|GYTYCTLMDGLCKE
ERIDEAVLLLDEMQSEGCSPS|PVIYNVLIDGLCKKGDLTRVTKLVDNMFLKGCVPN|EV
TYNTLIHGLCLKGKLDKAVSLLERMVSSKCIPN|DVTYGTLINGLVKQRRATDAVRLLSS
MEERGYHLN|QHIYSVLISGLFKEGKAEEAMSLWRKMAEKGCKPN|IVVYSVLVDGLCRE
GKPNEAKEILNRMIASGCLPN|AYTYSSLMKGFFKTGLCEEAVQVWKEMDKTGCSRN|KF
CYSVLIDGLCGVGRVKEAMMVWSKMLTIGIKPD|TVAYSSIIKGLCGIGSMDAALKLYHE
MLCQEEPKS|QPDVVTYNILLDGLCMQKDISRAVDLLNSMLDRGCDPD|VITCNTFLNTL
SEKSNSCDKGRSFLEELVVRLLKRQ|RVSGACTIVEVMLGKYLAPKTSTWAMIVREICKP
KKINAAIDKCWRNLCT

**b**

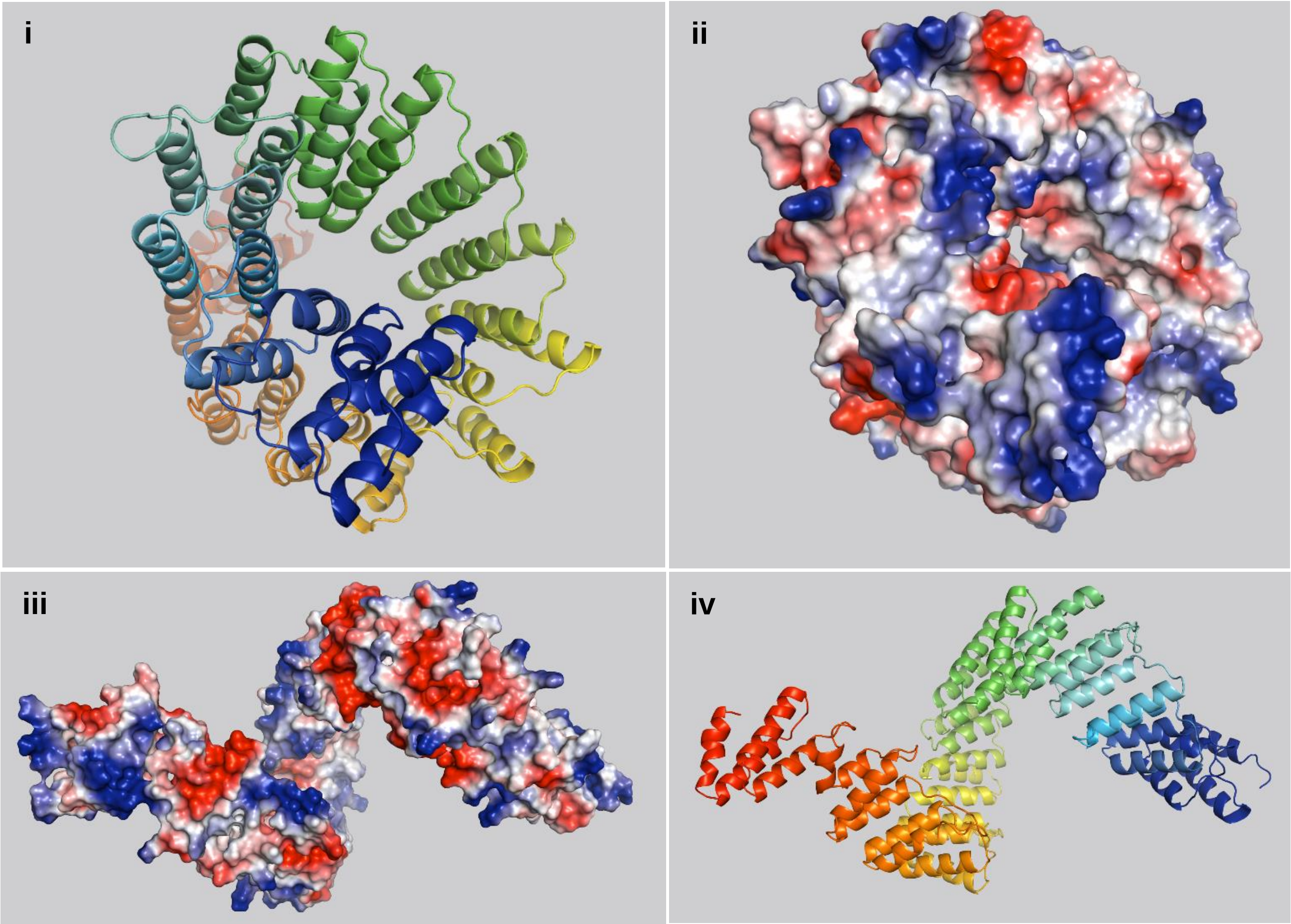


**Supplementary Figure S1. The topology of MSP1 protein.**

(a) The deduced amino acid sequence of MSP1 (At4g20090). The postulated regions, corresponding to the mitochondrial targeting sequence (36~37 amino acid long) is underlined and highlighted in magenta). The predicted cleavage sites of the Mitochondrial Processing Peptidase (MPP, 36 aa) or Intermediate Cleavage Peptidase 55 (ICP55, 37 aa) are indicated by box. The 15 PPR motifs (highlighted in blue) were predicted by the SMART (Letunic, *et al*. 2012), the Conserved Domain Database (CDD) (Marchler-Bauer, *et al*. 2003) and PPRCODE (Shen, *et al*. 2019) servers. (b) To get more of an insight on MSP1's mode of action, in particular of RNA recognition and binding, we performed an atomic model of the protein using the Phyre$^2$ server (Kelley and Sternberg 2009). The model structure of MSP1 (*i.e*. ribbon and surface views) were generated by the PyMol software suite (DeLano and Lam 2005). “i” and “iv” represent the ribbon structure from different angles, while panels “ii” and “iii” represent the hypothetical surface of MSP1 protein. The color code is red for negative values, white for near zero values, and blue for positive values.

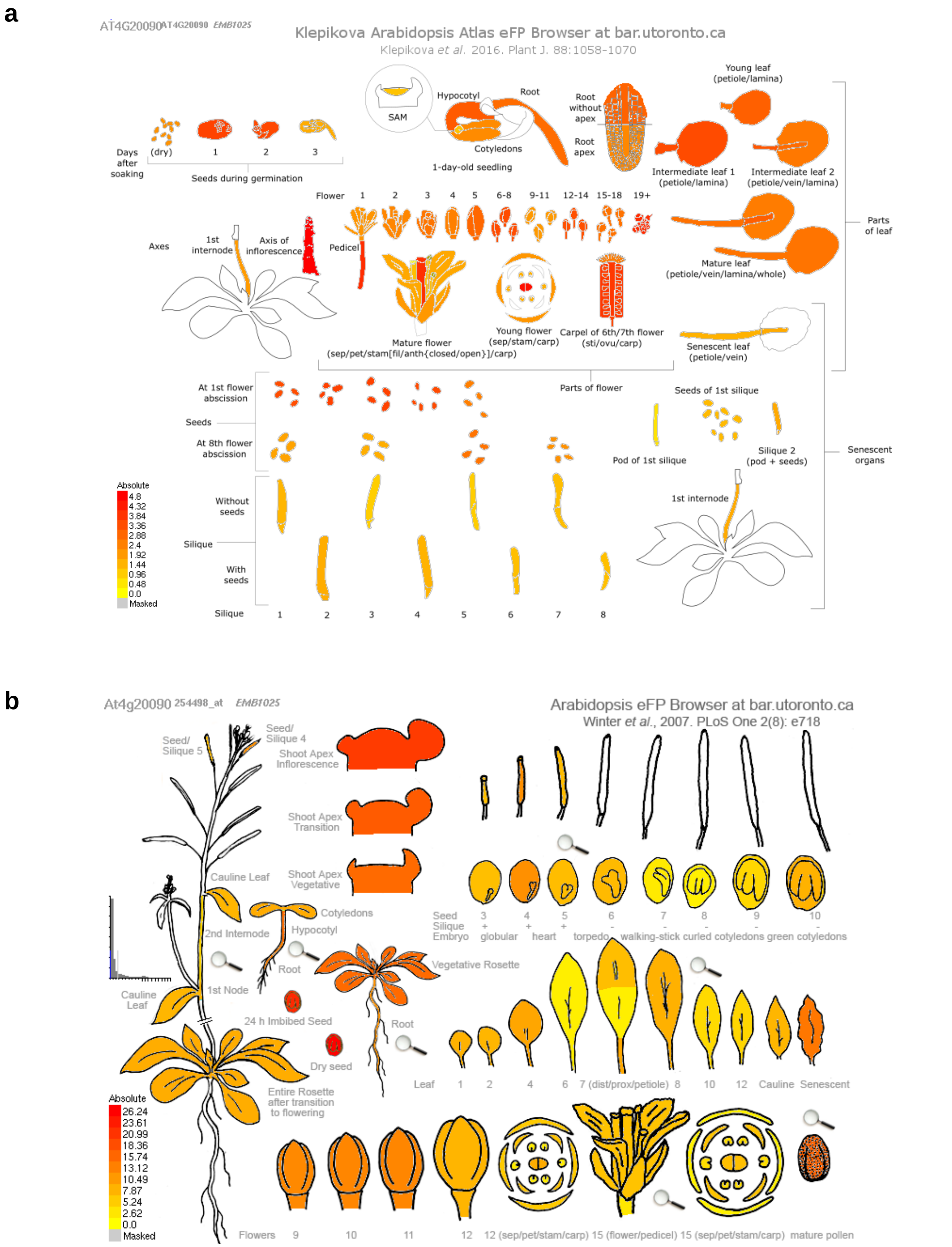


**Supplementary Figure S2. *MSP1* gene expression patterns at different tissues and during various developmental stages.**

The expression patterns of MSP1 were analyzed by publicly available microarray and high throughput sequencing databases, including The Arabidopsis Information Resource, TAIR; http://www.arabidopsis.org) and Genevestigator analysis toolbox (Hruz *et al.* 2008, Zimmermann *et al.* 2004).

5’…tatatagacatataaattagtatatataaatttgtttacttttaagattttctattttttagagaataaaaagttaaa
ttttgaatgacacatgtaaacatctgatcgtttgatttgacacgtggcacaatcctatgagttagcaattttgaaaccatata
ttttatataataagatccccaaaattattatgtatatgagattatttaccatttggatttgattacataaattttataaattaa
aaaaattattggtaaatgaattagaaaatttcacttttaatagtaattaagaagtaattaatcaattaattgagaggaaacaa
a **(SALK_018927)** aaaattgaaatttattgataactcttagtggcgaagaagagaagaagcagaaat**{SALK_018927_LB}**
caaatgtcatcgtttatcaacaaacctctaaattgatcgaatatcttcttcaatcttcatctctctttcaaatctgcaaatgag
agagctgtaagagttgcatcatctctatctctttctctctctcttgcgaaaacaccaaacttttgaaatccccaaataaggg
tttaagcataaaccctacgagttc**ATG**CCCAAATGCCCAATCCCAATCCGTATCAGCTTCTTCAGCTACTTTCTCAAAGAAAGT
CGAATCCTTTCGAGTAACCCAGTTAACTTCTCTATCCATCTTCGCTTCTCTTCTTCTGTTTCGGTTTCTCCTAACCCGTCAATG
GAAGTGGTGGAGAATCCATTGGAAGCTCCAATCTCGGAGAAGATGTTTAAATCAGCTCCCAAAATGGGTTCTTTCAAGTTGGGT
GATTCTACTTTATCTTCAATGATTGAGAGTTACGCCAATTCGGGTGATTTTGATTCGGTAGAGAAGCTTTTGAGTCGAATTAGA
**(SALK_142675)** TTAGAGAACAGAGTGATTATAGAGCGTAGTTTCATTGTTGTCTTTAGAGCTTATGGGAAAGCACATTTGCC
TGATAAAGCTGTAGACTTGTTTCATAGAATGGTTGATGAGTTTCGATGTAAACGTAGTGTTAAGTCGTTTAACTCTGTGTTGAA
TGTGATTATAAACGAGGGTCTTTATCATCGAGGATTGGAGTTTTATGATTATGTTGTGAATTCCAACATGAACATGAACATTTC
TCCTAATGGGTTGAGTTTTAATTTGGTCATTAAGGCTCTTTGTAAGTTAAGATTTGTGGATAGAGCTATTGAGGTCTTCAGAGG
AATGCCTGAGAGAAAGTGTTTGCCTGATGGTTATACGTATTGTACATTGATGGATGGGTTGTGTAAGGAGGAAAGGATTGATGA
GGCGGTTTTACTGTTGGATGAGATGCAGAGCGAGGGGTGTTCACCGAGTCCGGTTATATATAATGTGTTGATTGATGGATTGTG
CAAGAAAGGGGATTTGACACGGGTGACAAAGCTTGTGGATAATATGTTTCTTAAGGGATGTGTTCCTAATGAGGTTACTTATAA
TACGCTTATTCATGGTCTATGTCTTAAGGGTAAGTTAGATAAAGCTGTTAGTCTCTTGGAGCGGATGGTGTCTAGCAAATGTAT
CCCAAATGATGTCACATATGGAACGCTGATTAATGGGTTGGTTAAGCAAAGACGAGCAACGGATGCGGTTAGGTTGTTGAGCTC
TATGGAAGAAAGAGGGTATCATTTGAATCAGCATATTTATTCGGTCCTTATAAGTGGGTTGTTCAAAGAGGGAAAGGCCGAGGA
GGCTATGTCCTTGTGGAGGAAAATGGCAGAGAAAGGATGCAAGCCCAATATTGTTGTGTACAGTGTTCTTGTAGATGGTTTATG
CCGCGAAGGGAAACCAAATGAAGCTAAAGAGATCCTTAATAGAATGATTGCTAGTGGCTGCTTGCCCAATGCGTATACGTATAG
CTCATTGATGAAAGGATTCTTCAAAACTGGTCTTTGTGAAGAAGCGGTTCAAGTGTGGAAAGAGATGGATAAAACCGGATGTTC
TCGAAATAAGTTTTGTTATAGTGTTCTAATCGATGGTCTTTGTGGGGTTGGGAGAGTTAAGGAGGCTATGATGGTGTGGTCAAA
GATGCTCACTATCGGGATCAAACCTGATACAGTAGCTTATAGCTCAATTATAAAAGGTCTTTGTGGTATTGGCTCGATGGATGC
GGCCTTAAAACTTTACCATGAGATGTTATGCCAAGAAGAACCCAAGTCACAACCTGATGTTGTTACTTATAATATACTTCTTGA
TGGCTTATGTATGCAAAAAGATATCTCTAGAGCAGTAGATCTCTTGAACTCTATGTTAGATAGAGGTTGTGATCCCGATGTTAT
TACGTGTAACACCTTTTTGAATACTTTGAGCGAGAAGTCAAATTCTTGTGACAAAGGGAGGAGTTTTCTAGAAGAGCTTGTTGT
AAGGCTCTTAAAGCGTCAGAGAGTATCAGGCGCGTGTACGATTGTGGAAGTAATGCTGGGTAAGTATCTGGCACCGAAGACCTC
CACTTGGGCGATGATTGTTCGAGAGATCTGTAAACCAAAGAAGATCAATGCCGCCATTGATAAATGCTGGAGGAACCTGTGTAC
T**TGA**acactacagagcatttttgctcagatcccgtttccaccttaagagtctccatgcgtgggaacttttttgcgcttctggagt
tttggaataccagaaatgcgcgcgaaaaacaggtagtcagataaactatttttgaaattctaactttttaaacatgttacttga
tttatgcgcgagtaagaccacattagctaaggtttataaggatatgaaaatcccattttgttgctatt**(SALK_070654)**gaa
tcctccattgttaagatatgttctgtttcagacatgtttgatttgactatttgagatctttgtt…3’

**Figure S3. The nucleotide sequence of MSP1 gene and the precise locations of each of the T-DNA-insertional sites.**

The nucleotides sequence of *MSP1*, encoded by At4g20090 gene-locus. Underlined letters indicate to the 5’ and 3’ untranslated regions (UTRs), as indicated by the RACE analysis and TAIR database, while uppercased letters represent the open reading frame of mTERF22. The position of T-DNA insertions in *msp1-1* (SALK-142675), *msp1-2* (SALK-018927) and *msp1-3* (SALK- 070654).

**Supplementary Table S1.** Prediction of MSP1 PPR motifs and analysis of their RNA-binding code

**PPR motifs and PPR codes: undefined**

| Motif Start | Motif End | Motif Sequence | Fifth amino | Last amino : | PPR code | RNA base | Motif Lengt | ProSite Score |
|---|---|---|---|---|---|---|---|---|
| 40 | 74 | DSTLSSMIESYANSGDFDSVEKLLSRIRLENRVII | S | I | SI | ? | 35 | 7.552 |
| 75 | 109 | ERSFIVVFRAYGKAHLPDKAVDLFHRMVDEFRCKR | I | R | IR | ? | 35 | 6.993 |
| 111 | 145 | VKSFNSVLNVIINEGLYHRGLEFYDYVVNSNMNMN | N | N | NN | C>U | 35 | 6.982 |
| 150 | 184 | GLSFNLVIKALCKLRFVDRAIEVFRGMPERKCLPD | N | D | ND | U>C>G | 35 | 10.983 |
| 185 | 219 | GYTYCTLMDGLCKEERIDEAVLLLDEMQSEGCSPS | C | S | CS | A | 35 | 13.285 |
| 220 | 254 | PVIYNVLIDGLCKKGDLTRVTKLVDNMFLKGCVPN | N | N | NN | C>U | 35 | 11.488 |
| 255 | 289 | EVTYNTLIHGLCLKGKLDKAVSLLERMVSSKCIPN | N | N | NN | C>U | 35 | 13.329 |
| 290 | 324 | DVTYGTLINGLVKQRRATDAVRLLSSMEERGYHLN | G | N | GN | A>C | 35 | 10.731 |
| 325 | 359 | QHIYSVLISGLFKEGKAEEAMSLWRKMAEKGCKPN | S | N | SN | A | 35 | 13.066 |
| 360 | 394 | IVVYSVLVDGLCREGKPNEAKEILNRMIASGCLPN | S | N | SN | A | 35 | 13.176 |
| 395 | 429 | AYTYSSLMKGFFKTGLCEEAVQVWKEMDKTGCSRN | S | N | SN | A | 35 | 11.860 |
| 430 | 464 | KFCYSVLIDGLCGVGRVKEAMMVWSKMLTIGIKPD | S | D | SD | G>>C | 35 | 11.805 |
| 465 | 499 | TVAYSSIIKGLCGIGSMDAALKLYHEMLCQEEPKS | S | S | SS | A | 35 | 9.843 |
| 503 | 537 | VVTYNILLDGLCMQKDISRAVDLLNSMLDRGCDPD | N | D | ND | U>C>G | 35 | 12.649 |
| 538 | 573 | VITCNTFLNTLSEKSNSCDKGRSFLEELVVRLLKRQ | N | Q | NQ | ? | 36 | 6.522 |

**The RNA predicted by this PPR sequence:**
**(?) (?) (C>U) (U>C>G) (A) (C>U) (C>U) (A>C) (A) (A) (A) (G>>C) (A) (U>C>G) (?)**

**Notes:**

'-' represents no correlated RNA bases for this PPR code identified by biochemical assays (EMSA, ITC);

?' represents unknown bases, as no biochemical assays have been performed to identify the correlated RNA bases for the PPR code.

**Supplementary Table S2.** Oligonucleotides used in screening of individual T-DNA insertion lines in Arabidopsis and cloning of the *MSP1-GFP* gene-fusion construct.

| **Gene target** | **Gene I.D.** | **oligo name** | **Oligonucleotide sequence (5'-to-3')** |
|---|---|---|---|
| *PMS1* | At4g20090 | MSP1-F1 | CTCCCAAAATGGGTTCTTTC |
| | | MSP1-F2 | GAATGACACATGTAAACATCTG |
| | | MSP1-F3 | TCAATTAATTGAGAGGAAACAAA |
| | | MSP1-R1 | CATCAGGCAAACACTTTCTCT |
| actin2/8 | At3g18780 | Act2-F | TCTTCCGCTCTTTCTTTCCAAG |
| | | Act2-R | CTGGCGTACAAGGAGAGAAC |
| SALK T-DNA right border | pROK2 T-DNA | RB-10064 | CCAGATCCGGTGCAGATTATTTG |
| SALK T-DNA left border | pROK2 T-DNA | LB-6443 | CATCGCCCTGATAGACGGTT |
| SALK T-DNA left border | pROK2 T-DNA | LBb1.3 | ATTTTGCCGATTTCGGAAC |
| MSP1-GFP fusion | pSAT6-eGFP-N1 cloning | PMS1-GFP-F | ACATAGCCATGGACATGCCCAAATGCCCAATCCC |
| | | PMS1-GFP-R | CCATCAGGATCCCTCTAAAGACAACAATGAAACTA |
| MSP1-GFP fusion expression | pCAMBIA Cloning | PF_SAT6-EcoRI | ACATAGGAATTCTTCCCAGTCACGACGTTGTA |
| | | PR_SAT6-SmaI | ACATAGCCCGGGCAGCTATGACCATGATTACTG |

**Supplementary Table S3.** Lists of oligonucleotides used for the analysis of the splicing profiles of wild-type and mutant plants by RT-qPCR experiments.

| Gene | Forward primer | genome position | Reverse primer | genome position |
|---|---|---|---|---|
| *rpl2* | CCGAAGACGGATCAAGGTAA | 155542..155561 | CGCAATTCATCACCATTTTG | (157249..157268) |
| *rpl2* intron exon2 | TTAGGAAGAGCCGTACGAGG | 157142..157161 | CGCAATTCATCACCATTTTG | (157249..157268) |
| *rps3* | AGCCGAAGGTGAGTCTCGTA | 26990..27009 | CCGATTTCGGTAAGACTTGG | (28696..28715) |
| *rps3* intron1 exon2 | AGCCGAAGGTGAGTCTCGTA | 26990..27009 | TCTACGGCGGGGTCACTAT | (27088..27106) |
| *cox2* | TGGGGGATTAATTGATTGGA | 40518..40537 | TGATGCTGTACCTGGTCGTT | (42016..42035) |
| *cox2* intron1 exon2 | TGGGGGATTAATTGATTGGA | 40518..40537 | AGCAGTACGAGCTGAAAGGC | (40640..40659) |
| *ccmFc* | GTGGGTCCATGTAAATGATCG | 51765..51785 | CACATGGAGGAGTGTGCATC | (52856..52875) |
| *ccmFc* intron1 exon1 | CCCGGATCGAATCAGAGTT | 52747..52765 | CACATGGAGGAGTGTGCATC | (52856..52875) |
| *nad1* exon1-2 | GACCAATAGATACTTCATAAGAGACCA | 289014..289040 | TTGCCATATCTTCGCTAGGTG | (318039..318059) |
| *nad1* intron1 exon2 | GACCAATAGATACTTCATAAGAGACCA | 289014..289040 | CGTGCTCGTACGGTTCATAG | (289133..289152) |
| *nad1* exon2-3 | ATTCAGCTTCCGCTTCTGG | 287943..287961 | TCTGCAGCTCAAATGGTCTC | (289034..289053) |
| *nad1* intron2 exon2 | GGTTGGGTTAGGGGAACATC | 288936..288955 | TCTGCAGCTCAAATGGTCTC | (289034..289053) |
| *nad1* exon3-4 | AAAAGAGCAGACCCCATTGA | 147026..147045 | TCCGTTTGATCTCCCAGAAG | (287955..287974) |
| *nad1* intron3 exon4 | AAAAGAGCAGACCCCATTGA | 147026..147045 | GGGAGCTGTATGAGCGGTAA | (147113..147132) |
| *nad1* exon4-5 | AGCCCGGGATCTTCTTGA | 143401..143418 | TCTTCAATGGGGTCTGCTC | (147030..147048) |
| *nad1* intron4 exon5 | AGCCCGGGATCTTCTTGA | 143401..143418 | ACGGAGCTGCATCCCTACT | (143482..143500) |
| *nad2* exon1-2 | GCGAGCAGAAGCAAGGTTAT | 80109..80128 | GGATCCTCCCACACATGTTC | (81259..81278) |
| *nad2* intron1 exon2 | GCGAGCAGAAGCAAGGTTAT | 80109..80128 | CCCATTCCTAACCAGTGGAG | (80272..80291) |
| *nad2* exon2-3 | AAAGGAACTGCAGTGATCTTGA | 332947..332968 | AATATTTGATCTTAGGTGCATT TTC | (79761..79785) |
| *nad2* intron2 exon2 | CCCGATCCGATAGTTTACAA | 79641..79660 | AATATTTGATCTTAGGTGCATT TTC | (79761..79785) |
| *nad2* exon3-4 | GCGCAATAGAAAGGAATGCT | 330240..330259 | CTATGGGTCTACTGGAGCTAC CC | (333071..333093) |
| *nad2* intron3 exon4 | GCGCAATAGAAAGGAATGCT | 330240..330259 | GGCGAATTTCAAACTTGTGG | (330377..330396) |
| *nad2* exon4-5 | CAAAGGAGAGGGGTATAGCAA | 327932..327952 | TATTTGTTCTTCGCCGCTTT | (329793..329812) |
| *nad2* intron4exon4 | CTTATTCGTGGCAACCTTCC | 329705..329724 | TATTTGTTCTTCGCCGCTTT | (329793..329812) |
| *nad4* exon1-2 | ATTCTATGTTTTTCCCGAAAGC | 162115..162136 | GAAAAACTGATATGCTGCCTT G | (163664..163685) |
| *nad4* intron1 exon2 | CCGTATGATGCGGAAGTCTC | 163512..163531 | GAAAAACTGATATGCTGCCTT G | (163664..163685) |
| *nad4* exon2-3 | AATACCCATGTTTCCCGAAG | 163967..163986 | TGCTACCTCCAATTCCCTGT | (167228..167247) |
| *nad4* intron2 exon3 | GCGGAACGACCAGAAAAATA | 167110..167129 | TGCTACCTCCAATTCCCTGT | (167228..167247) |
| *nad4* exon3-4 | TTCCTCCATAAATTCTCCGATT | 167577..167598 | TGAAATTTGCCATGTTGCAC | (169651..169670) |
| *nad4* intron3 exon4 | TCTAGCTTGGTTCGGAGAGC | 169498..169517 | TGAAATTTGCCATGTTGCAC | complement(169651..169670) |
| *nad5* exon1-2 | TGGACCAAGCTACTTATGGATG | 141880..141901 | CCATGGATCTCATCGGAAAT | complement(142793..142812) |
| *nad5* intron1 exon2 | TGGACCAAGCTACTTATGGATG | 141880..141901 | TTCGCAAATAGGTCCGACT | (141962..141980) |
| *nad5* exon2-3 | TACCTAAACCAATCATCATATC | 190740..1907610 | CTGGCTCTCGGGAGTCTCTT | (140743..140762) |
| nad5 intron2-exon2 | GTACGATCGTGTCGGGTGA | 140656..140674 | CTGGCTCTCGGGAGTCTCTT | (140743..140762) |
| *nad5* exon3-4 | AACTCGGATTCGGCAAGAA | 22023..22041 | GATATGATGATTGGTTTAGGT A | (190740..1907610) |
| nad5 intron3-exon4 | AACTCGGATTCGGCAAGAA | 22023..22041 | GCCGTGTAATAGGCGACCA | (22150..22168) |
| *nad5* exon4-5 | AACATTGCAAAGGCATAATGA | 20695..20715 | GTTCCTGCGTTTCGGATATG | (21808..21827) |
| *nad5* intron4 exon5 | AACATTGCAAAGGCATAATGA | 20695..20715 | CCTGTAAACCCCCATGATGT | (20827..20846) |
| *nad7* exon1-2 | ACCTCAACATCCTGCTGCTC | 132118..132137 | AAGGTAAAGCTTGAAGATAAG TTTTGT | (133203..133229) |

| | | | | |
|---|---|---|---|---|
| *nad7* intron1 exon2 | ACGGTTTTTAGGGGGATCTG | 133128..133147 | AAGGTAAAGCTTGAAGATAAG TTTTGT | (133203..133229) |
| *nad7* exon2-3 | GAGGGACTGAGAAATTAATAGAGTACA | 133179..133205 | TGGTACCTCGCAATTCAAAA | (134362..134381) |
| *nad7* intron2 exon3 | AGTGGGAGAGCCGTGTTATG | 134182..134201 | TGGTACCTCGCAATTCAAAA | (134362..134381) |
| *nad7* exon3-4 | ACTGTCACTGCACAGCAAGC | 134718..134737 | CATTGCACAATGATCCGAAG | (135963..135982) |
| *nad7* intron3 exon4 | TAAAGTGAAGTGGTGGGCCT | 135797..135816 | CATTGCACAATGATCCGAAG | (135963..135982) |
| *nad7* exon4-5 | GATCAAAGCCGATGATCGTAA | 136007..136027 | AGGTGCTTCAACTGCGGTAT | (137951..137970) |
| *nad7* intron4 exon5 | CGGCCAAATGACTACAGGAT | 137856..137875 | AGGTGCTTCAACTGCGGTAT | (137951..137970) |
| rpl5B | CAGAAGACCTTTCCGTGCTC | chr5 15904477..15904496 | CACTGGAAACCGTGTGTTTG | (15904525..15904 544) |
| *actin2* | GGTAACATTGTGCTCAGTGGTGG | chr3 6476501..6476523 | AACGACCTTAATCTTCATGCTG C | (6476608..647658 6) |
| *matR* | AATTTTTGCGAGAGCTGGAA | 231141.. 231160 | TTGAACCCCGTCCTGTAGAC | (231313 .. 231294) |
| 18S nuclear rRNA | AAACGGCTACCACATCCAAG | chr3 14198079..14198098 | ACTCGAAAGAGCCCGGTATT | (14198179..14198 160) |
| *26S* rRNA | GACGAGACTTTCGCCTTTTG | 38542 … 38523 | CTTGGAGCGAATTGGATGAT | (38387… 38406) |

**Supplementary Table S4.** List of antibodies used for the analysis of wild-type and *msp1* mutants.

| Antibody | Protein I.D. | origin | serum | dilution | Reference / source |
|---|---|---|---|---|---|
| AtpB | Mitochondrial ATP-synthase subunit β | *Zea mays* | Mouse (monoclonal) | 1/5,000 | (Michael *et al.* 1993) |
| CA2 | γ-carbonic anhydrase-like subunit 2 | *Arabidopsis thaliana* | Rabbit (polyclonal) | 1/1,000 | (Perales *et al.* 2005, Sunderhaus *et al.* 2006) |
| Cox2 | Cytochrome oxidase subunit-2 | *Arabidopsis thaliana* | Rabbit (polyclonal) | 1/5,000 | Agrisera antibodies, AS04 053A |
| Nad9 | NADH-dehydrogenase complex subunit-9 | *Triticum spp.* | Rabbit (polyclonal) | 1/50,000 | (Lamattina *et al.* 1993) |
| RISP | Rieske iron-sulfur protein | *Arabidopsis thaliana* | Rabbit (polyclonal) | 1/5,000 | Gift of Prof. Ian Small, UWA |
